# Strengths and potential pitfalls of hay-transfer for ecological restoration revealed by RAD-seq analysis in floodplain *Arabis* species

**DOI:** 10.1101/474858

**Authors:** Hannes Dittberner, Christian Becker, Wen-Biao Jiao, Korbinian Schneeberger, Norbert Hölzel, Aurélien Tellier, Juliette de Meaux

**Affiliations:** Institute of Botany, University of Cologne, 50674 Cologne, Germany; Cologne Center for Genomics, University of Cologne, 50931 Cologne, Germany; Max-Planck-Institute for Plant Breeding Research, 50829 Cologne, Germany; Institute of Landscape Ecology, University of Münster, 48149 Münster, Germany; Center of Life and Food Sciences Weihenstephan, Technical University of Munich, 85354, Freising, Germany

**Keywords:** genetic diversity, RAD-seq, restoration genetics, hybridization, population structure, reference genome

## Abstract

Achieving high intraspecific genetic diversity is a critical goal in ecological restoration as it increases the adaptive potential and long-term resilience of populations. Thus, we investigated genetic diversity within and between pristine sites in a fossil floodplain and compared it to sites restored by hay-transfer between 1997 and 2014. RAD-seq genotyping revealed that the stenoecious flood-plain species *Arabis nemorensis* is co-occurring with individuals that, based on ploidy, ITS-sequencing and morphology, probably belong to the close relative *Arabis sagittata*, which has a documented preference for dry calcareous grasslands but has not been reported in floodplain meadows. We show that hay-transfer maintains genetic diversity for both species. Additionally, in *A. sagittata*, transfer from multiple genetically isolated pristine sites resulted in restored sites with increased diversity and admixed local genotypes. In *A. nemorensis,* transfer did not create novel admixture dynamics because genetic diversity between pristine sites was less differentiated. Thus, the effects of hay-transfer on genetic diversity also depend on the genetic makeup of the donor communities of each species, especially when local material is mixed. Our results demonstrate the efficiency of hay-transfer for habitat restoration and emphasize the importance of pre-restoration characterization of micro-geographic patterns of intraspecific diversity of the community to guarantee that restoration practices reach their goal, i.e. maximize the adaptive potential of the entire restored plant community. Overlooking these patterns may alter the balance between species in the community. Additionally, our comparison of summary statistics obtained from *de novo* and reference-based RAD-seq pipelines shows that the genomic impact of restoration can be reliably monitored in species lacking prior genomic knowledge.

## Introduction

Habitat degradation is an ever growing problem in our modern world, causing unprecedented loss of biodiversity and essential ecosystem services (Baillie, Hilton-Taylor, & Stuart, 2004). Thus, there is a growing demand for ecological restoration, i.e. measures assisting the recovery of ecosystems that have been degraded, damaged or destroyed (Society for Ecological Restoration International Science & Policy Working Group, 2004). However, ecological restoration is a difficult process rarely leading to full ecosystem recovery (Benayas, Newton, Diaz, & Bullock, 2009). Thus, the young field of restoration ecology, which studies ecological processes in the light of restoration, is vital to improve restoration practices (Bullock, Aronson, Newton, Pywell, & Rey-Benayas, 2011; Roberts, Stone, & Sugden, 2009; Suding, 2011).

One of the main goals of ecological restoration is the recovery of biodiversity, including both species richness and intraspecific genetic diversity (henceforth called genetic diversity). Genetic diversity has generally positive effects on ecosystems (reviewed in Hughes, Inouye, Johnson, Underwood, & Vellend, 2008). For example, experimentally increasing the genetic diversity of *Solidago altissima* increased primary above-ground biomass productivity and arthropod diversity (Crutsinger et al., 2006). Genetic diversity also boosts resistance of populations to invasion and environmental fluctuations, presumably because it enhances the adaptive potential of populations (Reed & Frankham, 2003; Vrijenhoek, 1994). For example, high diversity experimental *Arabidopsis thaliana* populations showed higher resistance against invasion by *Senecio vulgaris* than low diversity populations (Scheepens, Rauschkolb, Ziegler, Schroth, & Bossdorf, 2017). Moreover, *Zoestra marina* populations with higher genetic diversity showed increased biomass production, plant density and faunal abundance during an extremely warm period (Reusch, Ehlers, Hämmerli, & Worm, 2005). Concordantly, restored populations of *Z. marina* with increased genetic diversity showed longer plant survival, grew more rapidly and provided enhanced ecosystem services, measured by increased primary productivity, invertebrate density and nitrogen retention. This effect was stable in a range of environmental conditions along a water-depth gradient (Reynolds, McGlathery, & Waycott, 2012). These examples demonstrate the importance of genetic diversity for ecosystem function and stability and hence the need to consider population genetics in the design and evaluation of restoration practices, which is the focus of restoration genetics.

Restoration genetics can not only inform the planning of restoration efforts, e.g. by identifying suitable source populations, but also help evaluating the success of restoration projects, e.g. by monitoring genetic diversity in restored populations (Mijangos, Pacioni, Spencer, & Craig, 2015; Williams, Nevill, & Krauss, 2014). In fact, studies comparing the level of genetic diversity in pristine and restored populations frequently report limited success, with a reduction of genetic diversity in restored populations. This decline in genetic diversity may be caused by genetic bottlenecks in plant nurseries, biases introduced by seed harvesting strategies, founder effects during recolonization and/or unreliable commercial seeds (reviewed in Mijangos, Pacioni, Spencer, & Craig, 2015). By contrast, the transfer of seed-containing hay from pristine (donor) to restoration (donee) sites, termed hay-transfer, is expected to limit the loss of genetic diversity and maintain site-specific local adaption (Hufford & Mazer, 2003; Kiehl, Kirmer, & Shaw, 2014). In addition, this method has the unique feature that it can, theoretically, restore an entire community without altering the genetic composition of populations and thus is the best method available for restoring entire ecosystems (Hölzel & Otte, 2003; Kiehl, Kirmer, Donath, Rasran, & Hölzel, 2010). So far, however, there is no empirical support for the efficiency of this practice (Bucharova et al., 2017), especially since many species maintain seed banks in the soil. Indeed, the genetic diversity specific to the seed bank will not be sampled with the hay, although it is known that it can contribute significantly to the maintenance of diversity (Tellier, Laurent, Lainer, Pavlidis, & Stephan, 2011).

The field of restoration genetics has witnessed a major technological shift over recent years. Restoration genetics studies have initially relied on microsatellites and AFLP markers and thus provided a limited overview on patterns of genetic variation within and between restored or pristine populations (reviewed in Mijangos et al., 2015). Now, genotyping-by-sequencing (GBS) methods are beginning to be more broadly adopted (Gruenthal et al., 2014; Massatti, Doherty, & Wood, 2018; O’Leary, Hollenbeck, Vega, Gold, & Portnoy, 2018; Torres-Martinez & Emery, 2016). These methods drastically reduce sequencing costs through strategies to sequence a reduced portion of the genome, e.g. Restriction site Associated DNA-sequencing (RAD-seq) (Elshire et al., 2011; Etter, Bassham, Hohenlohe, Johnson, & Cresko, 2011; Peterson, Weber, Kay, Fisher, & Hoekstra, 2012). In contrast to previous methods (AFLP, microsatellites), GBS approaches sample proportions of the genome that are sufficiently large to allow resolving patterns of genetic diversity and spatial structure even at very local scale where overall levels of genetic diversity are low (Bradbury et al., 2015; Jeffries et al., 2016; Reitzel, Herrera, Layden, Martindale, & Shank, 2013). In principle, GBS approaches have a third major advantage: they are well suited to unravel genetic diversity in non-model species without prior genomic information. Yet, the accuracy of genotyping in the absence of a reliable reference genome has been questioned (Shafer et al., 2016). Since target species in restoration projects rarely coincide with species or genera with advanced prior genomic knowledge, it is important to assess whenever possible, whether conclusions from RAD-seq-based restoration genetics studies depend on the availability of a reference genome.

Flood-plain meadows are species-rich ecosystems, accommodating many endangered and stenoecious species adapted to the variable moisture regime. Due to increased agricultural land-use and river regulation a large proportion of flood-plain ecosystems are degraded and have become a target for ecological restoration. Here, we use RAD-seq to evaluate whether genetic diversity is maintained in flood-plain meadows restored by hay-transfer in the Upper Rhine valley in Germany (Hölzel & Otte, 2003, Donath, Bissels, Hölzel, & Otte, 2007). We focused on *Arabis nemorensis*, a stenoecious species typically restricted to flood-plain meadows, and thus strongly endangered in Central Europe (Schnittler & Günther, 1999). *A. nemorensis* is a short-lived, mostly biennial hemicryptophyte that is known to maintain a long-lived soil seed bank (Hölzel & Otte, 2004). It is part of the *Arabis hirsuta* species aggregate, which comprises several morphologically similar but ecologically diverse species, which diverged about 1.2 million years ago (Karl & Koch, 2014).

We performed RAD-seq for over 130 plants collected across pristine and restored sites. This dataset allows us to ask the following questions: i) what is the level of genetic diversity and structuration of the pristine sites that served as source populations for restoration? ii) do restored sites show a lower level of genetic diversity than the pristine sites? iii) how did restoration affect the distribution of diversity within and among restored sites? iv) is the use of a reference genome necessary to reliably characterize the impact of restoration on genetic diversity? This work demonstrates that a thorough genetic analysis of source and restored sites reveals the complex dynamics at stake in the restoration process. Through genetic analysis, we reveal the unanticipated co-occurrence of *A. nemorensis* with *A. sagittata,* a morphologically similar and closely related species from the *A. hirsuta* aggregate, which normally exhibits a preference for drier habitats such as calcareous grasslands (Hand & Gregor, 2006). We further show that restoration by hay-transfer has maintained and can even enhance local diversity. Our analysis further shows that the use of a reference genome yields higher estimates of genetic diversity, but does not affect the resolution of patterns of genetic variation within and between sites, indicating that this approach can find broad applications in the field.

## Methods

### Plant material and DNA extraction

The sampling area comprises the fossil, dyke-protected flood plain of the River Rhine near Riedstadt in Hessen, Germany. The area is dominated by arable fields, but also contains remnants of pristine flood meadow communities in low lying depressions that are submerged by ascending groundwater during high floods of the River Rhine. Since ca. 20 years, new flood meadow communities have been restored on ex-arable land using the transfer of green hay from the pristine sites sampled in this study as donors to overcome significant dispersal limitation (Hölzel & Otte, 2003). During this process, hay from different donor sites was placed in distinct yet adjacent patches, making admixture possible. Since restoration is still ongoing, restored sites differ in age (Table S1).

From previous monitoring of the sites we knew that populations of *Arabis* plants were not present every year in all sites. Thus, we sampled in two consecutive years to maximize the number of study sites. In the few sites sampled in both years, the genetic composition of the samples was not markedly different across years, so samples from both years were bulked for these sites (see Table S1, Figure S1). We harvested seeds from a total of 134 plants of *Arabis nemorensis/sagittata* in nine sites named from A to I, in order of collection. The presence of *A. sagitata* was not previously reported in these sites and was thus unexpected. Although some individuals showed the reduced stem leaf density and shorter siliques typical of *A. sagittata*, these phenotypic criteria were not always clearly distinguishable in the field, especially at the end of the season when siliques matured, so that the presence of the two species has remained overlooked in previous studies (Burmeier, Eckstein, Donath, & Otte, 2011). Species identity was thus determined by post-hoc analysis, after the RAD-seq analysis revealed the presence of two taxonomic units.

Four sites were sampled in pristine habitat (B, C, D and I) and five in restored habitat (A, E, F, G and H, Figure 1, Table S2). In site A two distinct stands of plants separated by about 100 m were sampled and were treated as sub-sites A-1 and A-2 throughout this study. To produce material for DNA extraction, we stratified seeds on wet filter paper for 6 days at 4°C in darkness. Afterwards we sowed seeds in soil (33% VM, 33% ED-73, 33% Seramis (clay granules)). After 4 to 6 weeks of growth in the greenhouse, we harvested about 200mg of leaf material from one offspring of each wild parent (genotype). We homogenized freshly harvested leaf material using a Precellys Evolution homogenizer (Bertin technologies) for 2×20 seconds at 6800 rpm. We extracted DNA using the NucleoSpin Plant II Mini kit (Macherey-Nagel) following the manufacturer’s instructions. We verified DNA quality using gel-electrophoresis with a 0.8% agarose gel. We measured DNA quantity using Qubit (broad-range kit) following manufacturer’s instructions.

**Figure 1:**
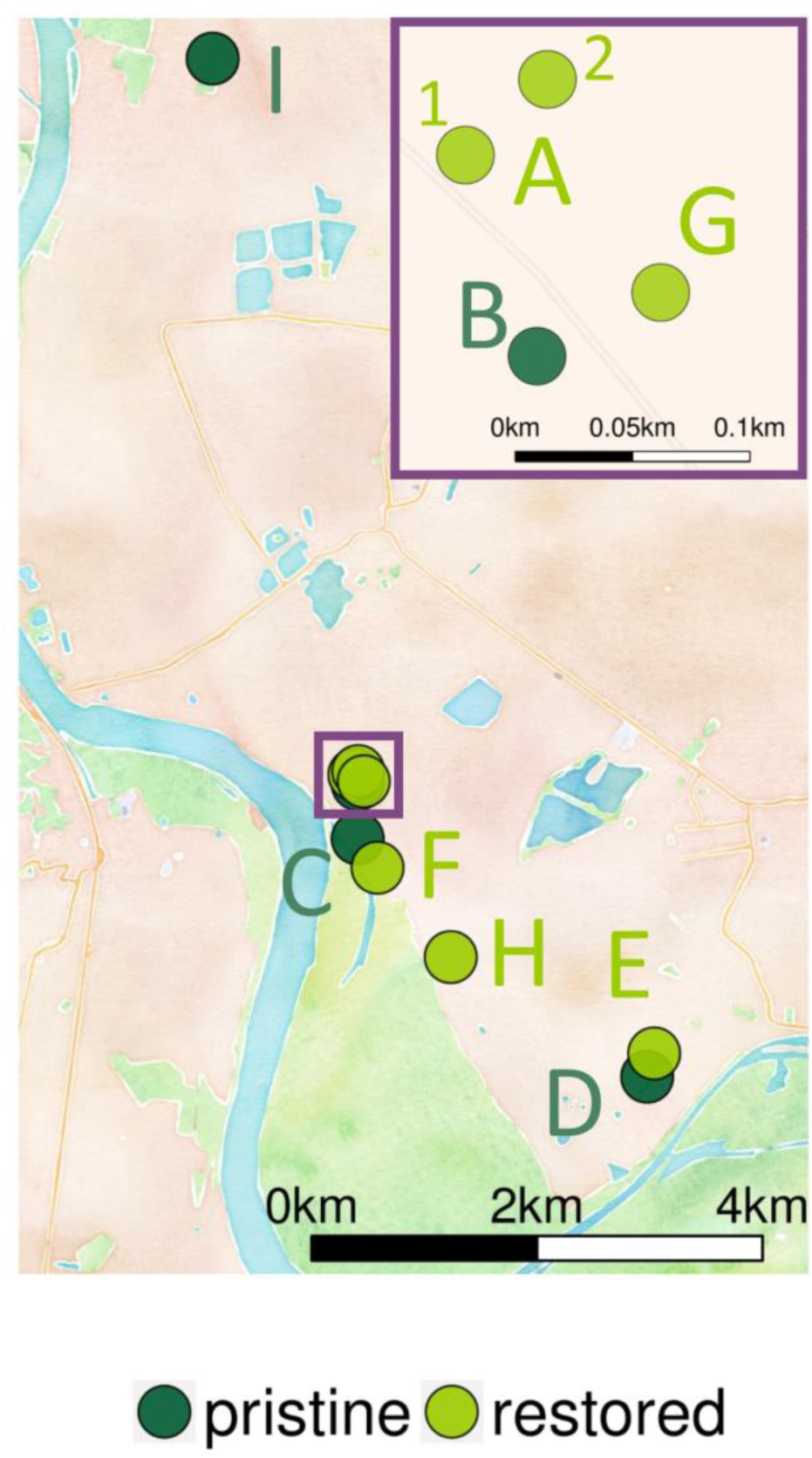
Overview map of studied sites. Each point on the map represents one site. The color of the points represents the type of the site, i.e. whether it is pristine or restored. The purple square represents the area shown in the zoomed inset. All sites, except site F are protected by a dyke.

### Draft genome assembly & annotation

To facilitate genotype calling we assembled a draft genome for one *A. nemorensis* accession (ID 29). Library preparation and sequencing was done at the Cologne Center for Genomics. Details about the sequencing, assembly and annotation of the genome are described in Document S1. Briefly, three libraries with insert sizes of 280 bp, 3 kbp and 6 kbp, respectively, were created and sequenced as 150 bp paired-end reads as part of an Illumina Hiseq 4000 lane for a total of 66 Gbp. We assembled reads using the *ALLPATHS-LG* assembler (Gnerre et al., 2011) with default settings. To further scaffold the genome, we generated 2.9 Gbp of PacBio sequence data. Library preparation and sequencing was done at the Max-Planck-Institute for Plant Breeding Research (Cologne, Germany). We scaffolded the genome using *OPERA-LG* with default settings (Gao, Bertrand, Chia, & Nagarajan, 2016). To achieve chromosome-level assembly, we created a whole genome alignment with the *Arabis alpina* reference genome (Jiao et al., 2017; Willing et al., 2015) and performed synthenic path assembly using the tool *Synmap2* (Haug-Baltzell, Stephens, Davey, Scheidegger, & Lyons, 2017) available at the *CoGe* website (Lyons & Freeling, 2008). The pseudo-chromosomes had a total size of 192 Mbp and were used for all following analyses. The size of the genome was estimated by flow-cytometry, which was performed commercially at Plant Cytometry Services (Didam, Netherlands). The estimated genome size was 274 Mbp. Thus, the assembly size was 70% of the genome size.

To detect and annotate Transposable elements (TE) we used the softwares RepeatModeler (Smit & Hubley, 2008) and RepeatMasker (Smit, Hubley, & Green, 2013). Protein-coding genes were annotated by integrating predictions of *ab initio* gene annotation tools and alignments of homologous proteins. Three different tools including Augustus v3.2.3 (Stanke & Waack, 2003), GlimmerHMM v3.0 (Majoros, Pertea, & Salzberg, 2004) and SNAP v2013 (Korf, 2004) were used to predict the initial gene models. Protein sequences from *A. thaliana, A. lyrata* and *A. alpina* (Arabidopsis Genome Initiative, 2000; Hu et al., 2011; Willing et al., 2015) were aligned to the assembly by the tool Exonerate v2.2.0 (Slater & Birney, 2005). Then, the *ab initio* predictions and protein alignment hits were further combined to build the consensus gene models by the tool EVidenceModeler (EVM) v2012 (Haas et al., 2008). Finally, TE related genes in these models were annotated by checking the TE annotation, blastp (Altschul, Gish, Miller, Myers, & Lipman, 1990) alignments with Plant TE related proteins and blastp alignments with *A. thaliana* proteins.

### RAD-sequencing and SNP calling

We genotyped 134 samples using the original RAD-sequencing (RADseq) protocol (Etter et al., 2011), with the following modifications (Document S1): i) we used the enzyme KpnI-HF (New England Biolabs) for DNA digestion, ii) we ligated digested DNA with complementary adapters containing one of ten different barcodes and a stretch of five random nucleotides, used for post-hoc removal of PCR duplicates (Table S3), iii) we created 14 pools of 10 barcoded samples each in equal amounts, iv) we used indexed reverse primers for amplification, described in (Peterson, Weber, Kay, Fisher, & Hoekstra, 2012), to allow multiplexing of pools. Libraries were sequenced on two Illumina HiSeq 4000 lanes with 2×150bp.

We used *FastQC* (Andrews, 2010) to quality-check the resulting reads. We trimmed adapters and removed reads shorter than 100bp using *Cutadapt* (Martin, 2011). We removed PCR duplicates based on a 5 bp stretch of random nucleotides at the end of the adapter, using the *clone_filter* module of *Stacks* version 1.37 (Catchen, Hohenlohe, Bassham, Amores, & Cresko, 2013). We de-multiplexed samples using the *process_radtags* module from *Stacks*. We filtered reads with ambiguous barcodes (allowed distance 2) and cut-sites, reads with uncalled bases and low-quality reads (default threshold).

For reference-based genotyping, we mapped reads using *BWA* (Li & Durbin, 2009) with default settings. We filtered mapped reads using *SAMtools* (Li et al., 2009) and custom python scripts using the following criteria to remove reads: mapping quality < 30, number soft-clipped bases > 30; reads were unpaired; mates mapped on different chromosomes; mate mapping distance > 700. The proportion of mapped reads varied slightly among species, with a mean of 48% for *A. nemorensis*, 43% for hybrids and 42% for *A. sagittata* (Table S4). We called genotypes using *SAMtools mpileup* and *VarScan2* (Koboldt et al., 2012) with the following options: base quality > 20; re-calculation of base quality on the fly (-E option); read depth > 14; strand filter deactivated; SNP calling p-value < 0.01. We filtered genotyped loci using *VCFtools* (Danecek et al., 2011) and custom python scripts removing loci with missing data in more than 5% of individuals and loci in masked (repetitive) regions. We clustered contiguous loci spaced less than 100 bp apart into RAD-regions. Regions with excessively low or high coverage are likely results of allele dropout or paralogous mapping, respectively. Thus, we removed regions fulfilling one of the following criteria: mean coverage of the region greater than twice the overall mean coverage; mean coverage of the region smaller than a third of the overall mean coverage; region maximum coverage greater than twice the mean maximal coverage over all regions; region shorter than 250bp; region longer than 1300bp.

After depth filtering, we still found loci with a high frequency of heterozygotes (up to 100%), which is unlikely in highly selfing species. Notably, in over 90% of these loci only one of the two homozygous genotypes was observed. Moreover, these highly heterozygous loci clustered in high density on single RAD-fragments. Thus, these loci likely resulted from paralogous mapping, which passed the depth filter due to sequencing depth variation among RAD-regions. Therefore, we removed RAD-regions containing loci with a frequency of heterozygotes greater than 20%. This threshold was picked to limit the impact of noise from mapping artifacts while still allowing for reasonable levels of heterozygosity, since some low-level outcrossing likely occurs. From the resulting genotype dataset, we extracted single nucleotide polymorphisms (SNPs) using *VCFtools* (Danecek et al., 2011).

For *de-novo*-genotyping we used the *Stacks* 2.2 *denovo_map.pl* pipeline (Catchen et al., 2013). The aim of the de-novo analysis was to test whether one would reach similar results and conclusions without the use of a reference genome. Thus, we performed this analysis without knowledge about the two species (since we gained this from the reference-based analysis) and ran *Stacks* for all samples combined. As recommended by the authors of the tool (Rochette & Catchen, 2017), we first used a subset of 15 representative genotypes to tune the parameters (-M and -n) of the algorithm, which control the number of mismatches between stacks within (M) and between (n) individuals. We varied M and n from 1 to 9. For each set of parameters, we analysed the number of loci shared between 80% of the samples. This measure peaked at the value six for M and n. Thus, we used this value for both parameters for the full analysis. We ran the *denovo_map.pl* pipeline using 0.01 as the p-value threshold for calling genotypes and SNPs, and otherwise default options. We used the *populations* program to create a VCF-file for further analysis using the following filters: 5% maximum missing data per locus; 20% maximum observed heterozygosity per locus; locus must be present in all sites.

### Population genetics statistics

We did all statistical analysis using R version 3.4.4 (R Development Core Team, 2008) and provide a supplemental R Markdown file (Document S2). The following packages were used for plotting: *ggplot2* (Wickham, 2009), *ggmap* (Kahle & Wickham, 2013), *ggthemes* (Arnold et al., 2017), *ggsn* (Baquero, 2017) and *heatmap3* (Zhao, Guo, Sheng, & Shyr, 2015). We performed all analysis for the reference-based and the *de-novo*-based dataset and compared the results. We used the *vcfR* package (Knaus & Grünwald, 2017) to load VCF-files into R and make the SNP data available for processing with other libraries. Based on our annotation we determined whether SNPs are in coding regions and whether they are synonymous or non-synonymous using the *PopGenome* package (Pfeifer, Wittelsbuerger, Li, & Handsaker, 2018). We performed principal component analysis (PCA) of SNP data for all samples using the *adegenet* package (Jombart et al., 2016). Missing data was scaled to the mean for PCA.

Based on the first principal component most individuals (83%) could be assigned to one of two distinct taxonomic groups. We used molecular methods to assign species labels to the two taxonomic groups. We sequenced the internal transcribed spacer (ITS) sequence of nine individuals, three from the first (left) and five from the second (right) cluster and one located between the clusters (Table S5). Primers for amplification were taken from (Mummenhoff, Franzke, & Koch, 1997). We used the sequences as input to the taxonomy tool of the *Brassibase* website (Kiefer et al., 2014). As a complementary approach, we downloaded the 612 bp long reference ITS sequences of all species in the *Arabis hirsuta* group from Brassibase (Kiefer et al., 2014). Using the software Seaview (Gouy, Guindon, & Gascuel, 2010), we aligned these sequences to the ITS sequences of our samples using the MUSCLE algorithm (Edgar, 2004) (Document S3) and created a phylogeny using PhyML (Guindon & Gascuel, 2003) with default settings. The ITS sequence does not allow distinction between *A. sagittata* and *A. hirsuta*. Yet, the species can be distinguished based on ploidy as *A. sagittata* is diploid and *A. hirsuta* tetraploid (Karl & Koch, 2014). Leaf samples were therefore collected from twelve individuals (seven putative *A. sagittata/A. hirsuta*, two putative *A. nemorensis*, three putative hybrids; Table S6) and their relative DNA content per cell (in comparison to a company-internal standard (*Vinca minor*)) determined by Plant Cytometry Services (Didam, Netherlands). Additionally, we included one sample from each of two independent *Arabis hirsuta* populations sampled in 2016 in Bavaria (see Table S2). We inferred the ploidy of our samples by comparing relative DNA content: individuals of the same ploidy should have similar DNA content, tetraploids should have twice the DNA content of diploids.

We conducted all population genetic analysis separately for the two species. Individuals, which were not assigned to any species, were likely interspecific hybrids and excluded from further analysis. We used the *pegas* library to calculate within-site genetic diversity (Nei’s π; average pairwise nucleotide differences) for each site (Paradis, Jombart, Schliep, Potts, & Winter, 2016), excluding sites with less than two individuals per respective species. To scale the estimates of π, we divided the average number of pairwise nucleotide differences among samples by the total number of successfully genotyped bases, excluding all bases which failed any of our previously described genotype filters (missing data, region heterozygosity, region depth and length). For the de-novo pipeline we extracted the total number of genotyped bases from *Stacks* output. We calculated correlation coefficients between reference-based and *de-novo*-based π estimates using Pearson’s method. We calculated pairwise F_ST_ (Nei, 1987) and genetic distance (Cavalli-Sforza & Edwards, 1967) between all pairs of sites using the *hierfstat* package (Goudet & Jombart, 2015). Negative F_ST_ values were set to zero. Differences of genetic distance and F_ST_ among pristine and restored sites were tested using a Wilcoxon-rank-sum-test on pairwise distance matrices. We tested for correlation between the distance matrices of the reference and *de novo* datasets using a Mantel test with 10000 permutations.

### Admixture analysis

For ADMIXTURE analysis (Alexander, Novembre, & Lange, 2009), we converted VCF-files to bed-files using PLINK (Purcell, 2009; Purcell et al., 2007). First, we conducted ADMIXTURE analysis for all samples combined for K=1 to K=10 (reference-pipeline only). Then for each of the species and pipeline (reference/*de novo*), we ran ADMIXTURE analysis for K=2 to K=10, with 10 iterations of cross-validation each. Before plotting, we normalized clusters across runs using CLUMPAK (Kopelman, Mayzel, Jakobsson, Rosenberg, & Mayrose, 2015). We created plots using a custom R-script.

## Results

### RAD-sequencing uncovers two hybridizing species

Unless otherwise stated, all described results were obtained using the reference-based pipeline. We genotyped 134 individuals from 10 sites – 4 pristine and 6 restored (Figure 1) – yielding 3.6 Mb of sequence of which 32,880 single nucleotide positions were polymorphic (SNPs). Only 20% of SNPs were in coding regions, 40% and 56% of which were synonymous and non-synonymous, respectively (4% unassigned). To visualize patterns of genetic diversity across sites, we conducted principal component analysis (PCA) for all individuals (Figure 2, top). Almost 80% of the total genetic variation was explained by the first principal component, which separated most individuals into two clearly defined clusters, likely representing taxonomic units. To determine species identity, we performed phylogenetic analysis of the ITS region of nine individuals (Table S5) using two different methods (see methods), which gave the same results. We confirmed that one taxonomic unit was *A. nemorensis* (Figure S2-3). The ITS region of individuals from the other taxonomic unit was identical to that of *A. sagittata* and *A. hirsuta,* a sequence that differs by at least 3 nucleotides from all other ITS sequences of known species of the complex (Figure S3-4). Yet, these sibling species can be distinguished based on ploidy, as *A. hirsuta* is tetraploid and *A. sagittata* diploid (Karl & Koch, 2014). Since all twelve tested samples from the Rhine populations had the same genome size, which was half of that of the two *A. hirsuta* samples (Table S6), we conclude that the second cluster most likely corresponds to the diploid species *A. sagittata*. Average genetic distance (d_XY_) between *A. nemorensis* and *A. sagittata* was 8.1e-03, i.e. 8 fixed SNP differences for 1000 bp.

**Figure 2:**
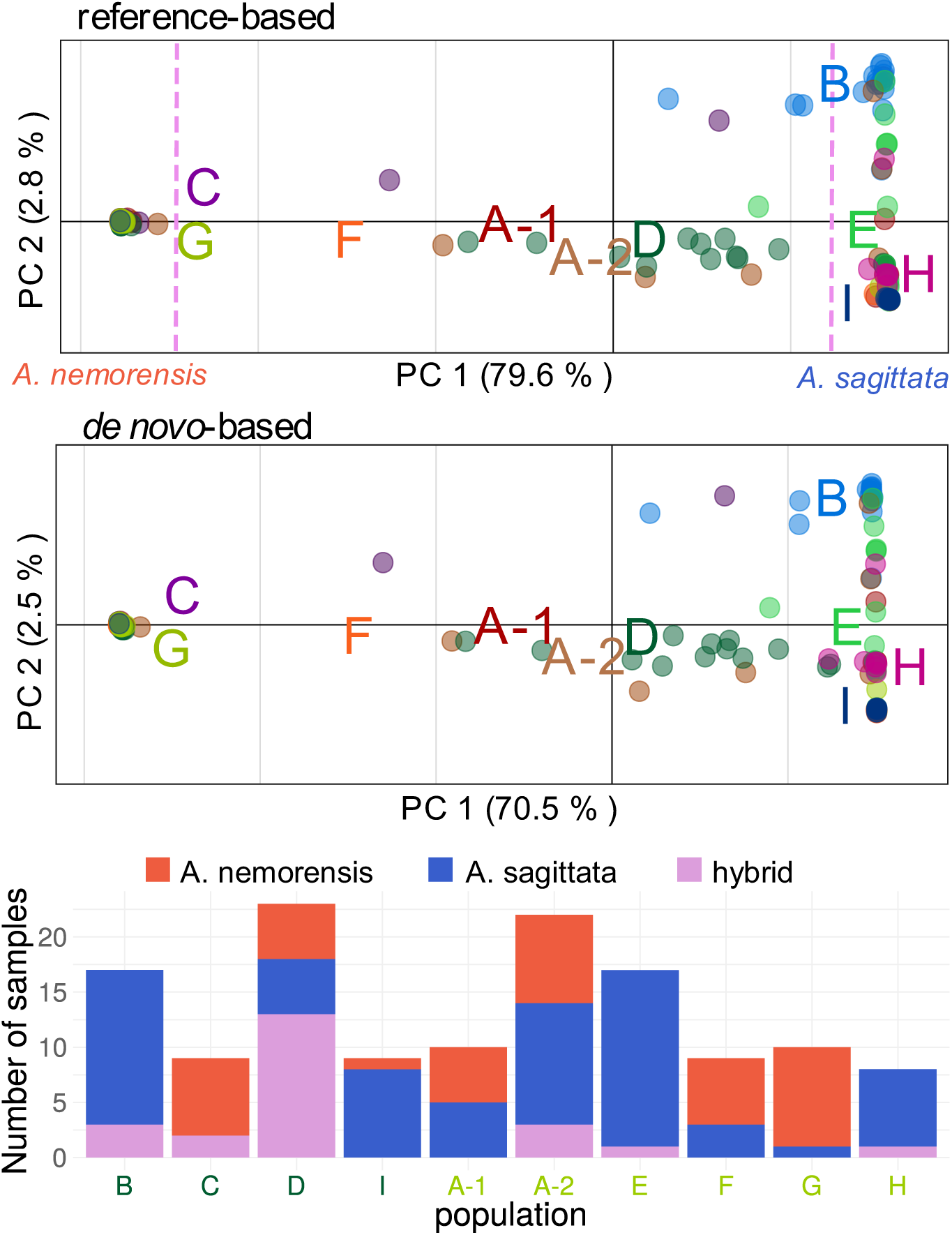
Presence of two hybridizing *Arabis* species in study sites. Top) Plot of the first two principal components (PC) of genetic variation based on the reference-based pipeline. Colors distinguish the different sites, which are labeled with letters of the corresponding color. Site labels are in the centroid of the respective site (with offset to avoid overlaps). Values in brackets in the axis labels show the amount of variance explained by each PC. Note that the first PC explains about 80% of the total variation and splits most individuals into two clearly defined clusters. The two clusters along PC1 correspond to two distinct species, identified here as *Arabis nemorensis (left)* and most probably *Arabis sagittata (right)*. Species identity is indicated by text labels below the plot. Points in between the two clusters are likely natural interspecific hybrids. The dashed lines show the thresholds used for distinction between species and hybrids throughout this study. Middle) Plot showing the same analysis as above but based on the de-novo-pipeline. Bottom) Stacked barplot showing the distribution of species among the study sites. Population labels are colored by type: darkgreen = pristine, lightgreen = restored. Species are distributed heterogeneously among sites.

Twenty-three individuals showed a positioning along the first PC that was intermediate between the two clusters, suggesting they were interspecific hybrids. Most of these hybrids were closer to *A. sagittata* on the first PC. Since F1-hybrids should be located exactly in the middle between the two species, this suggests that they are somewhat fertile and preferentially back-cross with *A. sagittata.* Laboratory observations have shown that hybrids are indeed fertile (Dittberner, personal observation). Additionally, in an ADMIXTURE analysis of all samples with K=2, most hybrids showed ancestry over 50% from *A. sagittata*, as expected from preferential back-crossing to this parent (Figure S5).

Overall our sample was composed of 31% *A. nemorensis*, 52% *A. sagittata* and 17% hybrids. The species composition differed among sites (Figure 2, bottom). *A. nemorensis* was present in 7 sites (3 pristine, 4 restored), *A. sagittata* in 9 sites (3 pristine, 6 restored) and hybrids in 6 sites (3 pristine, 3 restored). Notably, the pristine site D was dominated by hybrids with over 56%.

### No reduction of genetic diversity in restored sites

We computed species-specific estimates of genetic diversity within each site, excluding hybrid genotypes. The *A. nemorensis* dataset consisted of 2746 SNPs. Levels of genetic diversity (π) varied up to two-fold among sites, ranging from 6.6e-05 in A-2 to 1.4e-04 in A-1 (Figure 3, left; Table S7). Total diversity was 1.5e-04. However, pristine and restored sites did not differ significantly in their level of diversity (mean difference= +10% in restored; W = 4, p = 1).

**Figure 3:**
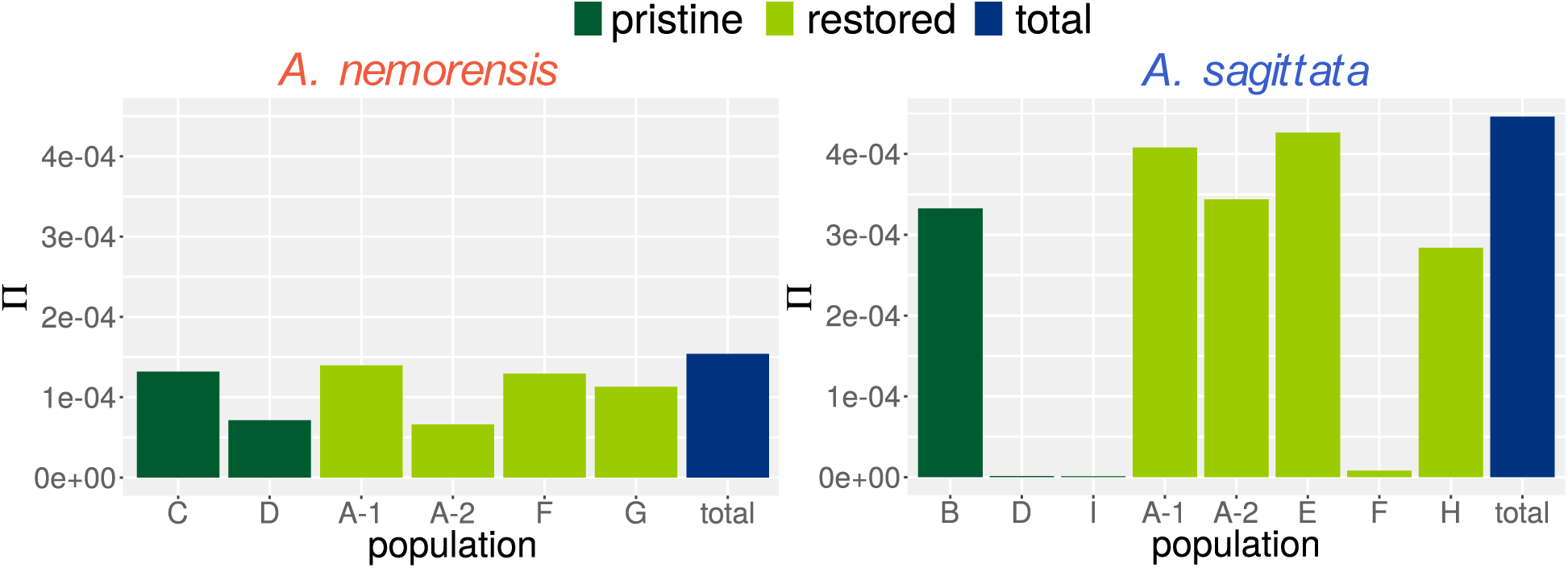
No reduction in genetic diversity in restored sites. Barplot of average pairwise genetic diversity (π) within each site of each species. Bar-color indicates the type of site.

The *A. sagittata* dataset consisted of 6366 SNPs. Total genetic diversity in *A. sagittata* was about three times as high as in *A. nemorensis*. Yet, in contrast to *A. nemorensis*, genetic diversity differed strongly among sites, ranging from 1.03e-06 in site I to 4.26e-04 in site E (Figure 3, right; Table S7). Notably, genetic diversity was low in two of three pristine sites. In contrast, we found high levels of diversity in all restored sites, except site F. However, the overall difference between pristine and restored sites was not significant (mean difference= +163% in restored sites; W=13, p=0.14).

Since hybridization potentially enables gene flow between the two species we also compared levels of genetic diversity for both species combined, including the hybrids. Overall genetic diversity increased by an order of magnitude and mixed sites were more diverse than mostly pure sites, as would be expected (Figure S6). Again, restored sites did not show significantly different levels of genetic diversity from pristine sites (mean difference= +22%; W=13, p=0.91).

### Restoration reduces population structure and facilitates recombination

To quantify the degree of population structure, we estimated genetic distance and differentiation (F_ST_) among all pairs of sites (Table S8). Genetic distance among *A. nemorensis* sites ranged from 0.03 to 0.31 and F_ST_ estimates from 0 to 0.5 (Figure 4, A+C). Population structure was slightly more pronounced among pristine sites than restored sites: Mean genetic distance was 0.26 among pristine sites and 0.13 among restored sites (W=17, p=0.047); mean F_ST_ was 0.37 among pristine sites and 0.25 among restored sites (W=14, p=0.26).

**Figure 4:**
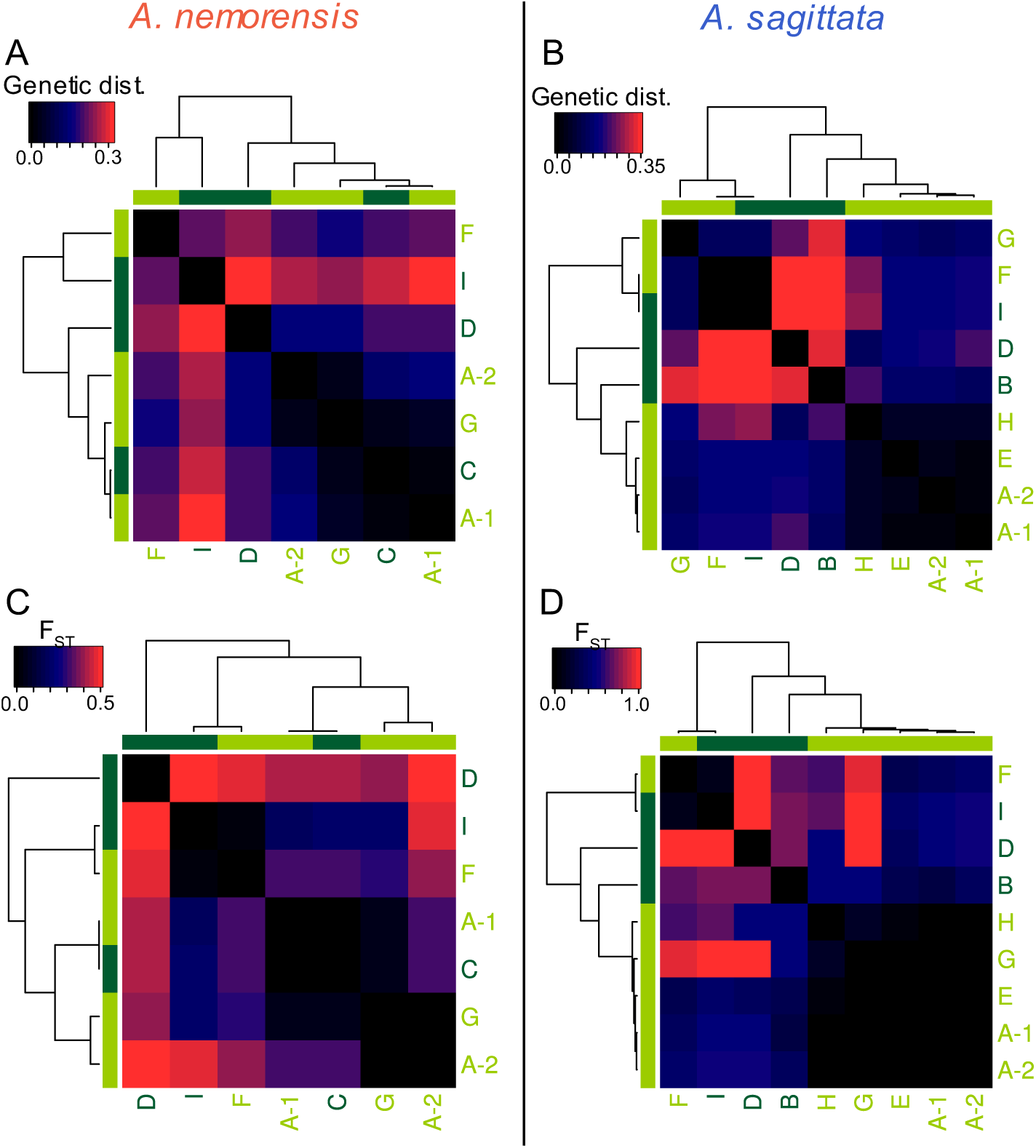
Overall strong population structure with signatures of admixture in restored sites. This panel shows different measurements of population structure of each species. A-D) Heatmaps showing pairwise comparisons between all sites of genetic distance (Cavalli-Sforza chord distance) and FST, respectively. Values of the respective variable are indicated by tile color, corresponding to the color scale in the top-left corner of each heatmap. Sites were clustered based on the respective variable, as indicated by the dendrograms. Colors of text labels and bars next to the grid indicate the type of the site (same color scheme as Figure 3).

In *A. sagittata*, genetic distance ranged from 0.32 to 0.34 and F_ST_ estimates from 0 to 0.91 (Figure 4, B+D). The difference in population structure between pristine and restored sites was stronger than in *A. nemorensis*: mean genetic distance was 0.33 among pristine sites and 0.12 among restored sites (W=45, p=0.002); mean F_ST_ was 0.81 among pristine sites and 0.2 among restored sites (W=43, p=0.015).

Reduced differentiation among restored sites suggests that genetic material of pristine sites was mixed in restored sites. To visualize this in more detail, we conducted ADMIXTURE analysis for each species. This analysis assumes a given number (K) of genetic clusters (ancestral populations) and assigns cluster ancestry proportions to each individual, allowing for mixed ancestry. For *A. sagittata*, K=6 was estimated as the optimal value (Figure 5). Up to K=4, the three pristine sites consisted of three distinct genetic clusters, which were mixed in all but one of the restored sites. For higher values of K, pristine population B consisted of two clusters, with pure and admixed individuals in equal proportions, in agreement with the higher genetic diversity observed at this site. All individuals from other pristine sites had pure ancestry from a single cluster per population.

**Figure 5:**
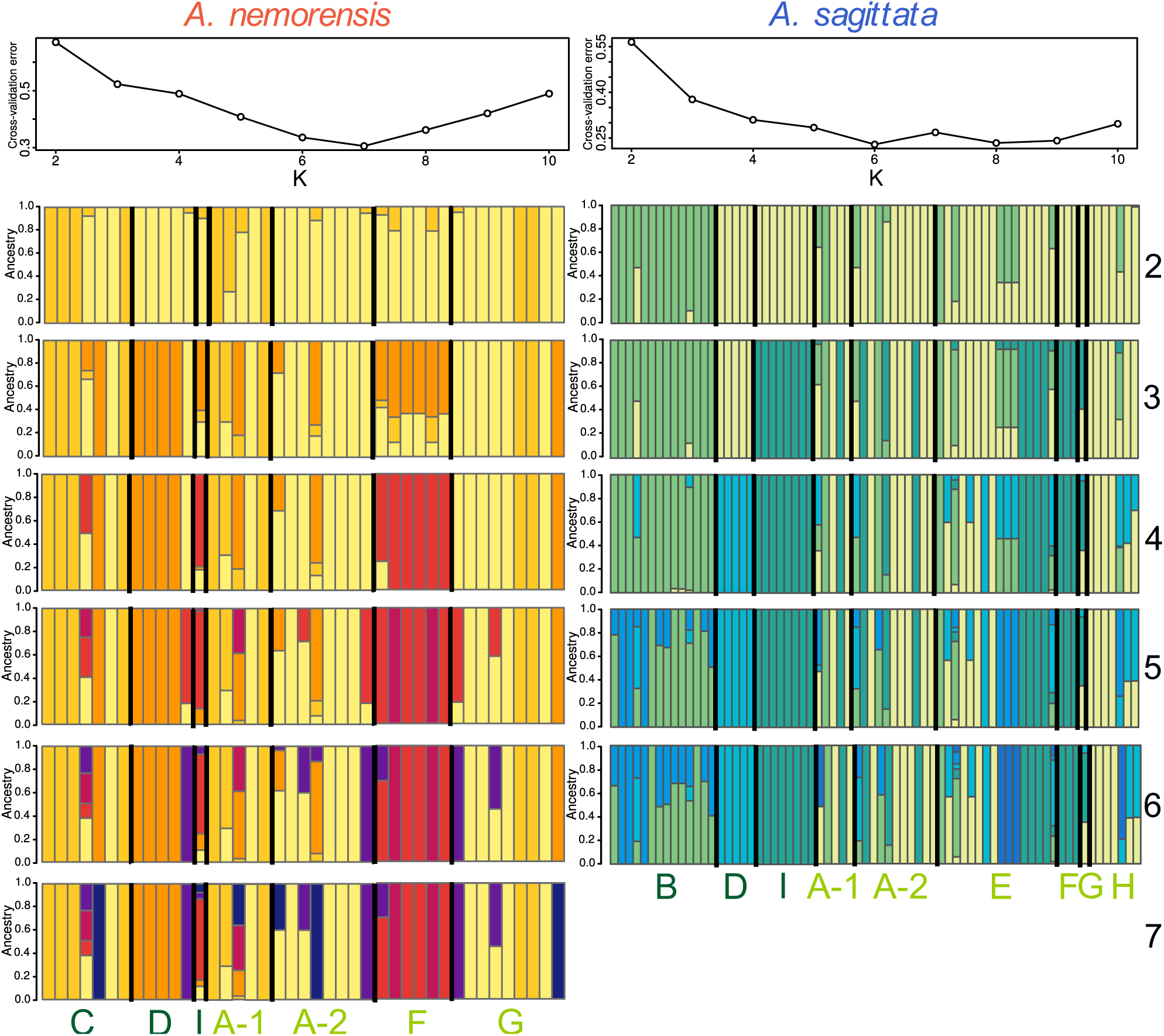
Increased impact of genetic admixture of pristine sites in *A. sagittata*. The figures shows a comparison of ADMIXTURE results for both species. The top panels show the cross-validation (CV) error for all tested values of K. The following panels show the ancestry proportions of each cluster for each individual for all values of K until the first minimum of the CV-error curve. The numbers on the right represent the value of K. Site labels are colored according to the type of the site: light-green = restored, dark-green = pristine.

For *A. nemorensis*, K=7 was estimated as the optimal value for clustering (Figure 5). In contrast to *A. sagittata*, we found individuals with different or mixed ancestry within populations of *A. nemorensis*, even at low values of K, indicating more genetic mixture than in *A. sagittata*. Since genetic structure was less pronounced for *A. nemorensis*, the restoration procedure did not impact the genetic distribution of this species, by contrast with *A. sagittata*.

### *De-novo*- and reference-based summary statistics reach the same conclusion

Finally, we tested whether a reference genome is required to determine the impact of restoration on genetic diversity by comparing results from a reference-based and a *de novo* pipeline. We found that estimates of genetic diversity, genetic distance or F_ST_ yielded by the two methods were highly correlated, with all correlation coefficients being greater than 0.95 (maximum p < 0.001, Figure 6). However, we observed that estimates of genetic diversity generated without a reference genome were deflated, especially for high diversity sites (Figure 6), by a median factor of 0.83 for *A. nemorensis* and 0.36 for *A. sagittata*. Estimates of genetic distance were underestimated by a median factor of 0.61 in both species. Interestingly, however, both pipelines yielded almost identical estimates of F_ST_ (median factor of 0.93) and revealed a very similar extent of admixture between sites (Figures S7-8). Moreover, the presence of the two species and their hybrids was detected with both methods (Figure 2 middle). We concluded that, in this study, the use of a reference genome was not required to determine the impact of restoration on genetic diversity. Since the two pipelines coincide, we could further conclude that the distribution of variation in two species and across pristine and restored sites reported above is not the result of possible mapping biases to the reference genome.

**Figure 6:**
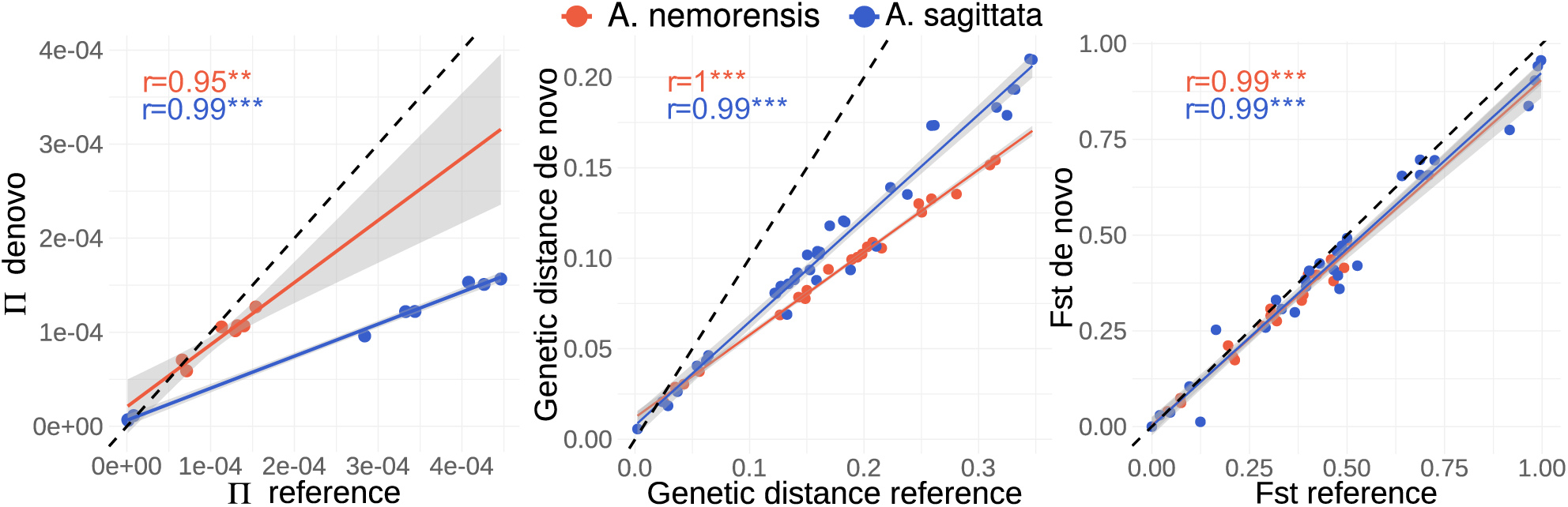
De-novo- and reference-based summary statistics are highly correlated. Plots show the correlation of summary statistics based on the reference-based and the de-novo-based pipeline. Line and dot colors represent the species. Each dot represents one individual. Lines represent a linear fit through the dots and the gray shadows indicate the error of the fit. The dashed line represents a hypothetical 1:1 relationship of the variables. Correlation-coefficients (Pearson’s product-moment correlation) are indicated as colored text corresponding to the species. Stars indicate the level of the coefficient: * * p< 0.01, * * * p<0.001.

## Discussion

### A reference-genome is not required to characterize the impact of restoration

While RAD-seq and related methods are a cheap tool to acquire genotype information across the genome without need for a reference genome (Elshire et al., 2011; Etter et al., 2011; Peterson et al., 2012), the reliability of *de novo* assembly pipelines has been questioned (Shafer et al., 2016). The availability of a reference genome allowed us to ask whether reference-based read mapping pipelines yielded distinct conclusions from a pipeline based on *de novo* read assembly. Conversely, this also allowed testing whether the use of an *A. nemorensis* reference genome to map *A. sagittata* samples could have biased our conclusions. The results from both pipelines were highly correlated. Thus, comparative analysis of sites was reliable with both pipelines. However, the *de-novo*-pipeline (*Stacks*, Catchen et al., 2013) underestimated the amount of genetic diversity compared to the reference-based pipeline. This is in contrast to a previous study comparing different RAD-seq pipelines (Shafer et al., 2016), where *Stacks* produced slightly inflated estimates compared to reference-based pipelines. However, in the same study, other de novo pipelines produced substantially lower estimates of genetic diversity (Shafer et al., 2016). Thus, the magnitude of genetic diversity estimates might depend on the study system and pipelines/parameters used, and caution is advised when comparing these estimates across studies.

In contrast, both pipelines agreed for analyses comparing diversity between species or sites (e.g. F_ST_, ADMIXTURE). Thus, we conclude that RAD-seq is an efficient tool to characterize the distribution of diversity, even in the absence of a reference genome. We therefore hope that this study will pave the way for exploring how species, with diverse life-history and uncharacterized genomes, will be maintained after hay transfer or how modalities of hay transfer affects not only single species but the balance between multiple species in the community.

### First documented presence of *A. sagittata* in floodplain meadow habitat

We detected two independent genetic clusters. Based on ITS sequences, ecological preference and morphological descriptors, the first one appears to represent *bona fide A. nemorensis* individuals. Taxonomic boundaries in the *Arabis hirsuta* clade can be tenuous, yet ITS sequence differences between the two clusters strongly suggests that they belong to distinct species (Karl & Koch, 2014). ITS sequences, ploidy level and morphological evidence collectively indicated that individuals in the second genetic cluster are likely to belong to *Arabis sagittata*. Species barriers are often incomplete, and patterns of divergence between closely related species of the same genus can be strongly intertwined. Population genomics approaches and wider sampling will be needed to consolidate our understanding of the *A. sagittata* taxonomic group and its historical relationship with the *A. nemorensis* taxon. It is indeed possible that gene flow between the species is ancient and not restricted to the Rhine population. Population genomics can indeed yield detailed view on the process and history of speciation in plant genera (see e.g. Novikova et al., 2016).

We did not anticipate the presence of the *A. sagittata* in flood-plain habitats because it is predominantly found in warm and dry habitats (Hand & Gregor, 2006). However, agricultural land-use and flood regulation have considerably modified the flood-plain ecosystem: flooding is contained by a dyke and the ground water level has decreased (Hölzel & Otte, 2001). Man-made modifications of the environment can change selection regimes and facilitate the establishment of non-native species (Byers, 2002; Crooks, Chang, & Ruiz, 2011; Fukasawa, Miyashita, Hashimoto, Tatara, & Abe, 2013; Tyrrell & Byers, 2007). Thus, it is possible that *A. sagittata* migrated into the flood-plain ecosystem after human river regulation decreased the frequency and severity of flooding. We want to stress that the species co-occurred both in pristine and restored sites. Thus, the contact between the species was not caused by the restoration efforts. In fact, we observed that *A. sagittata* is more frequent than *A. nemorensis* in our sample, which raises the concern that this species may be in the process of displacing *A. nemorensis*. This trend could be enhanced by climate change, which may lead to conditions that further favor the xero-thermophylic *A. sagittata*. Additionally, if *A. sagittata* is often mistaken for *A. nemorensis* in flora reports on flood-plain environments, the remaining *A. nemorensis* population in Central Europe might be even smaller than is currently assumed.

### The parallel transfer of two sympatric relatives shows that hay transfer maintains genetic diversity

Hay-transfer has proven to be a particularly successful method for establishing new populations of target species in ecological restoration across a variety of herbaceous vegetation types (Coiffait-Gombault, Buisson, & Dutoit, 2011; Hölzel & Otte, 2003; Kiehl et al., 2010; Kiehl & Wagner, 2006; Török et al., 2012). Furthermore, hay-transfer is often seen as the gold standard to preserve local levels of genetic diversity and adaptation (Vander Mijnsbrugge, Bischoff, & Smith, 2010). The latter is probably the main reason, why it is increasingly used in in ecological restoration (Kiehl et al., 2014). The aim of this study was to characterize the level of diversity in the pristine source sites and document the impact of hay-transfer on the genetic diversity in restored sites. Although our initial plan was to focus on *A. nemorensis*, a typical representative of species-rich floodplain meadows, the unanticipated presence of *A. sagittata* in our sample allowed us to compare the genetic effects of restoration by hay-transfer on the two species.

Several years after restoration, we did not find a significant difference in genetic diversity between pristine and restored sites for either of the two species. Thus, the hay-transfer method can restore populations with levels of diversity indistinguishable from the source populations in the long-term. Our findings are in agreement with studies on population life-stage structure and dynamics comparing pristine and restored sites of *A. nemorensis/A. sagittata* in the same region (Burmeier et al., 2011). This outcome is particularly remarkable given that *A. nemorensis* has a long-term seed bank, with up to 25,000 germinable seeds* m^-2^, which was not transferred to restored sites (Burmeier et al., 2011). Although species for which the genetic diversity present in the seed bank tends to differ more strongly from the above ground diversity may fare differently after hay-transfer, we note that populations restored with alternative methods, e.g. spontaneous recolonization (Vandepitte et al., 2012) or propagated seed mixtures (Espeland et al., 2017; Fant, Holmstrom, Sirkin, Etterson, & Masi, 2008), both excluded the seed bank and revealed a reduction of diversity (Mijangos et al., 2015). Thus, hay-transfer might be superior to other restoration methods not only in restoration success (Hölzel & Otte, 2003; Kiehl et al., 2010) but also in transferring genetic diversity.

### The modalities of habitat restoration can modify the relative adaptive potential of species in the ecosystem

Genetic diversity was low in *A. nemorensis* and *A. sagittata,* compared to other outcrossing or selfing Brassicaceae species (Mattila, Tyrmi, Pyhäjärvi, & Savolainen, 2017; Onge, Källman, Slotte, Lascoux, & Palmé, 2011). In fact, genetic distance between species was only on the order of magnitude as diversity found within populations of *Arabidopsis lyrata* (Mattila et al., 2017). Genetic diversity within our species was similar to that reported for *Arabis alpina* populations in Scandinavia (Laenen et al., 2018). As in our study, these populations are selfing and located on the margin of the species’ range (Jalas & Suominen, 1994), two factors often coinciding with lower levels of genetic diversity. Since populations with low genetic diversity may suffer from increased genetic load and decreased adaptive potential, the transfer of non-local material is often envisaged to preserve endangered species (Breed, Stead, Ottewell, Gardner, & Lowe, 2013; Weeks et al., 2011). This approach is however controversial. Strategies that introduce non-local seeds can decrease population fitness either by introducing maladapted genotypes (Crémieux, Bischoff, Müller-Schärer, & Steinger, 2010; McKay, Christian, Harrison, & Rice, 2005) or by causing outbreeding depression (Frankham et al., 2011). As a compromise, a strategy of mixing regional seeds was recently proposed (Bucharova et al., 2018). Our results show that this strategy was unintentionally implemented in the examined restoration effort: ADMXITURE analysis showed that pristine *A. sagittata* sites are dominated by one or two ancestral groups, with very low genetic diversity within each group. Yet, in some of the restored sites, admixture of low diversity *A. sagittata* groups took place, leading to a strong increase in genetic diversity and decrease in population structure. ADMIXTURE analysis also revealed that this led to genetic recombination between distinct haplotypes in *A. sagittata,* possibly increasing the adaptive potential of restored sites in this species. Increased genetic connectivity between populations has indeed been shown to help maintain or even increase genetic diversity in the long term (DiLeo, Rico, Boehmer, & Wagner, 2017). Fitness assays of recombined *A. sagittata* genotypes are needed to verify whether this local admixture has reinforced the establishment of this species in restored floodplain meadows.

Interestingly, we find less pronounced population structure in pristine sites of *A. nemorensis* than in *A. sagittata*, suggesting a comparatively higher level of gene flow. The restoration of this species is therefore less likely to benefit from post-transfer admixture. More so, our results suggest that it is possible that *A. nemorensis* is at increased disadvantage in restored habitats, if admixture and resulting recombination were to favor the emergence of more competitive genotypes in *A. sagittata*.

The coexistence of these species in the Rhine floodplain ecosystem will be further impacted by the ongoing hybridization dynamic that our analysis uncovers for the first time in this system and which occurred independently of the restoration effort. *A. nemorensis* could receive alleles conferring drought-adaptation from the xero-thermophilic *A. sagittata* that may enhance its ability to cope with increased drought exposure in its habitat. The genomic composition of the hybrids, however, also indicates that hybrids back-cross preferentially with *A. sagittata*. Gene-flow from *A. nemorensis* could facilitate adaptation of *A. sagittata* to the flood-plain environment. Such adaptive introgressions could also potentially accelerate the extinction of *A. nemorensis*. While hybridization is common in plants (Mallet, Besansky, & Hahn, 2016), well documented cases of adaptive introgression are rare and require elaborate experiments (Goulet, Roda, & Hopkins, 2017; Suarez-Gonzalez, Lexer, & Cronk, 2018). Such experiments are now warranted to determine the impact of hybridization in this sympatric species complex.

## Conclusions

Clearly, genetic analysis helps with species identification as sibling species can be difficult to distinguish morphologically even for specialists. A unique feature of hay-transfer restoration approaches is that the whole plant community can be transplanted. It is therefore particularly important to determine the composition of the source populations to limit the spread of non-target species (Bickford et al., 2007). Our study demonstrates that hay-transfer has maintained genetic diversity in restored populations. However, we also note that it may have inadvertently contributed to increase the genetic diversity and adaptive potential of only one of the two species, due to differences in the genetic makeup of the donor populations. This might lead to a competitive advantage of one over the other species in the long term, potentially disturbing the balance in the community. On the one hand this shows that restoration by hay-transfer may enhance the adaptive potential, which is especially important in the face of a rapid climate change. On the other hand, this also highlights that understanding the underlying genetics of the community to be transferred is a pre-requisite for the design of restoration strategies that promote the maintenance of both endangered ecosystems and endangered species.

## Acknowledgements

We thank Matthias Harnisch for providing information about the populations, Markus Koch for insightful discussions about the *A. hirsut*a tribe, Eric Schranz for his assistance with the pseudo-chromosome assembly and Gregor Schmitz for helpful feedback. Further, we thank Janine Altmüller and the team of the Cologne Center for Genomics for their assistance in RAD-seq optimization, library preparation and sequencing. This work was partly funded by the German Research Foundation ‘Deutsche Forschungsgemeinschaft DFG [DFG priority program 1529 ‘ADAPTOMICS’].

## Data availability

The genome assembly, annotation, raw reads, ITS sequences and RAD-seq samples will be uploaded to EMBL ENA upon acceptance (available upon request for review). VCF files and custom scripts will be uploaded to Dryad repository upon acceptance (available upon request for review). An R Markdown file describing the population genetic analysis is provided in the supplemental material.

## Author contributions

JdM, HD, AT and NH conceived the study. HD and NH collected plant material; HD and CB prepared material for sequencing; WBJ, KS and HD were responsible for bioinformatic processing. JdM, HD and NH analyzed data and wrote the manuscript with significant contributions from CB, WBJ, KS and AT.

## Notes

#### Summary of Updates

We added a more detailed description of phylogenetic analysis of ITS and ploidy analysis to identify species and discuss the results more carefully. We made minor changes in several parts of the manuscript and figures to improve clarity.

