## Supplemental Figures for "Strengths and potential pitfalls of hay-transfer for ecological restoration revealed by RAD-seq analysis in floodplain *Arabis* species"

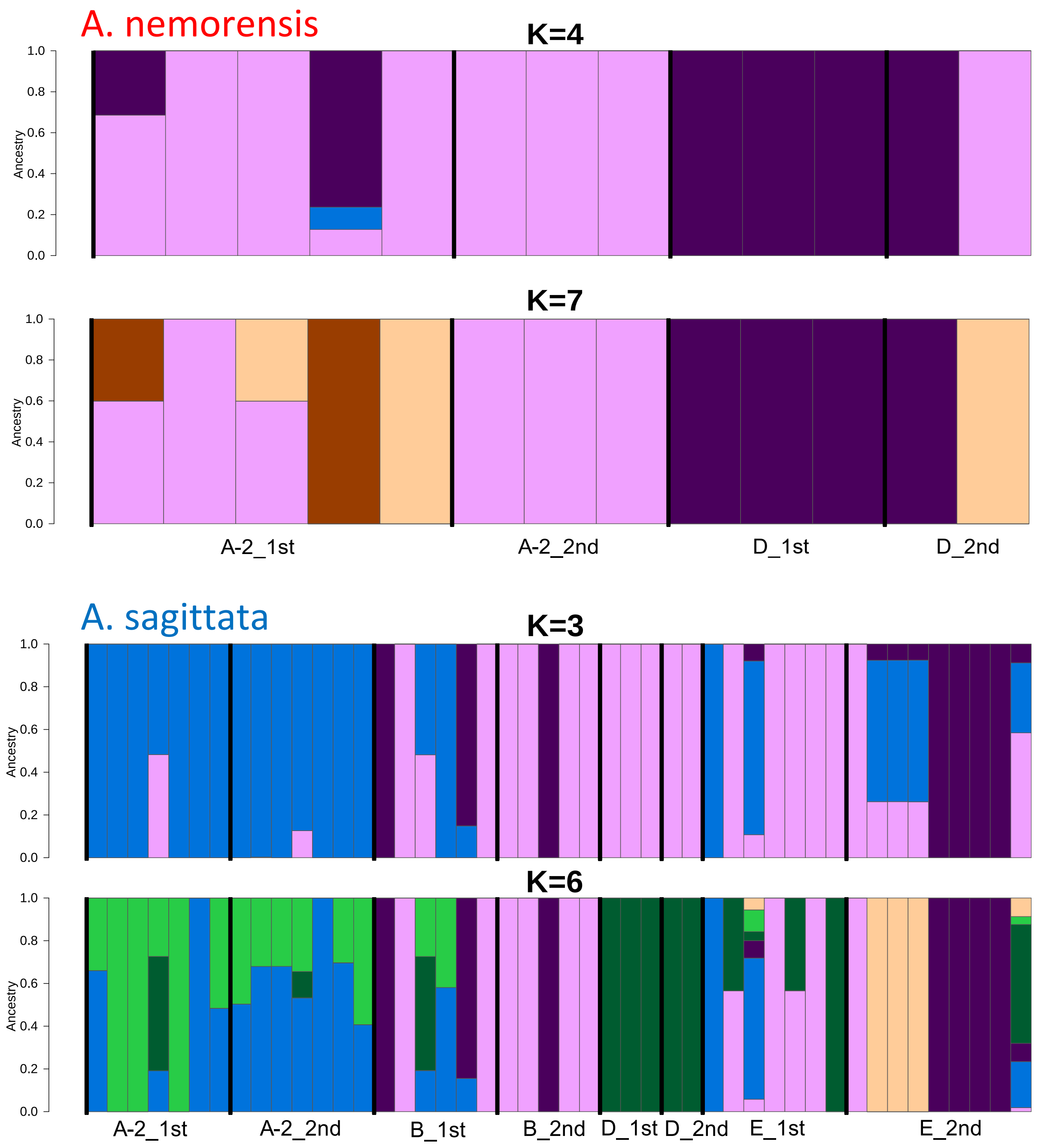


*Figure S 1: ADMXITURE plots of populations with two-year samples, split by year. Data was taken from the analysis for Figure 5 only rearranged to visualize differentiation among years. The lower K is the one used for the main figure; the higher K is the one with the lowest cross-validation error*

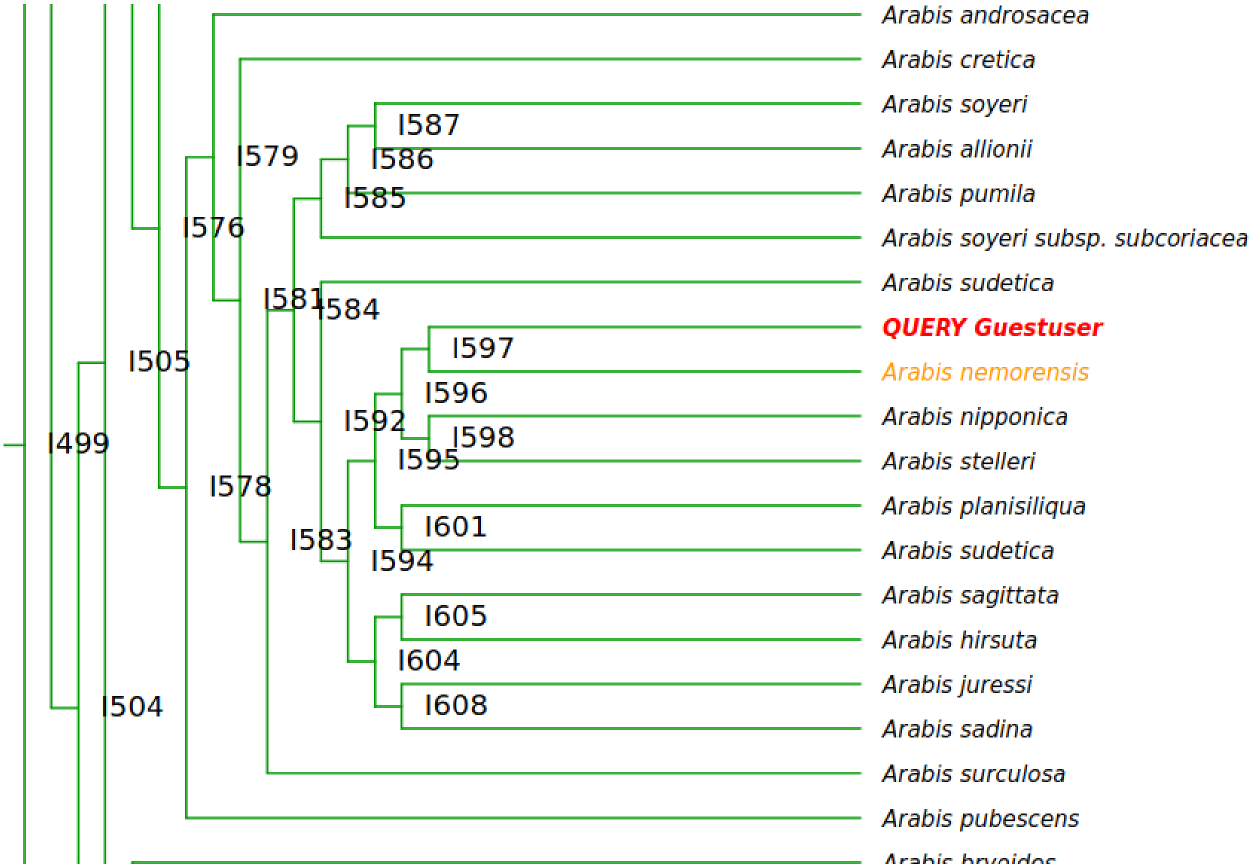


*Figure S 2: Example output from Brassibase for a sample of the first cluster (genotype ID 29). Query Guestuser represents the query sequence and the closest match is marked in orange.*

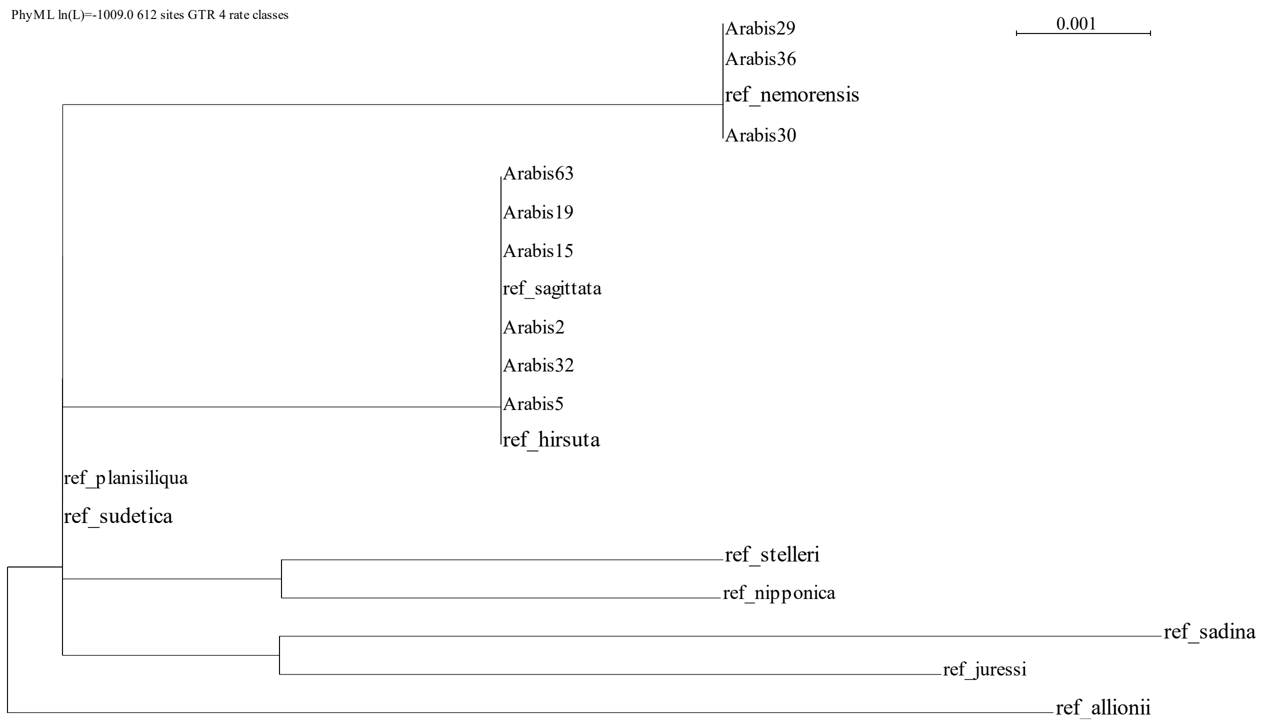


*Figure S 3: Phylogenetic tree based on the ITS sequence of nine samples of the Rhine population (marked as Arabis…) and reference sequences (marked as ref_...) downloaded from Brassibase.*

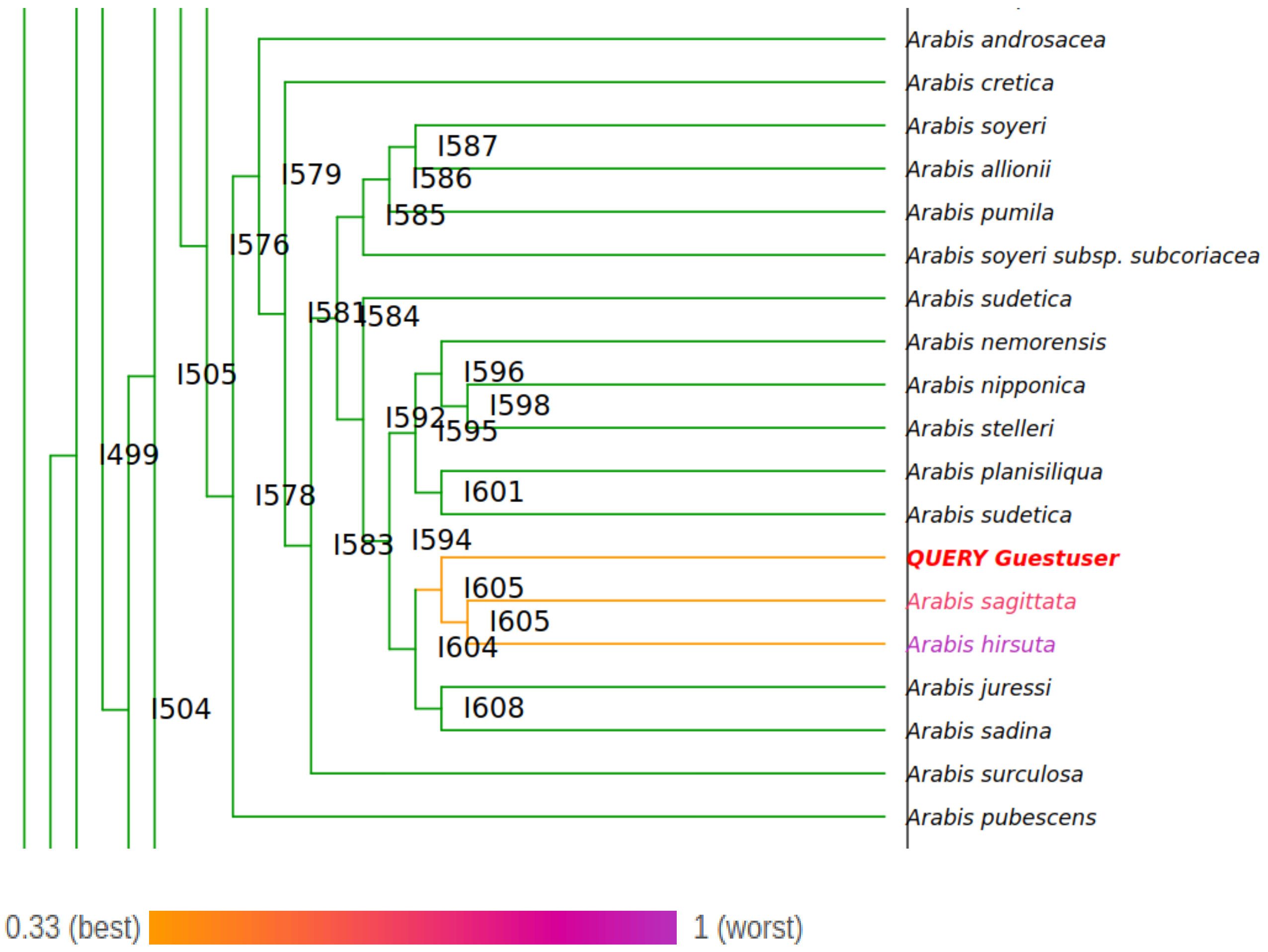


*Figure S 4: Example output from Brassibase for a sample of the second cluster (genotype ID 2). Query Guestuser represents the query sequence. Match likelihoods of the two closest matches are color-coded, as indicated by the color scale.*

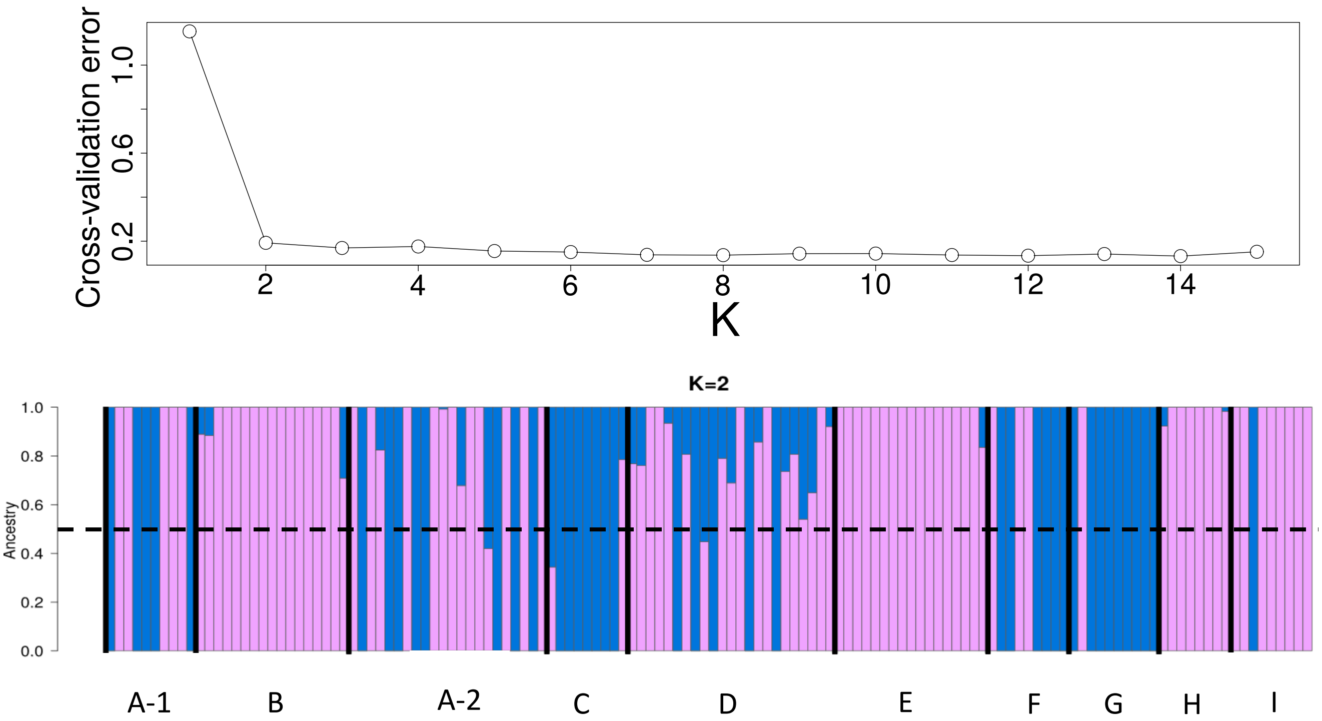


*Figure S 5: Top) Cross-validation error of ADMIXTURE analysis for different values of K. Bottom) ADMIXTURE plot for all individuals including hybrids for K=2. Blue represents the A. nemorensis cluster and pink the A. sagittata cluster. Individuals are ordered by site, sites are separated by black lines and site labels are below the plot. The dashed black line is approximately at ancestry level 0.5. This shows that most hybrids have A. sagittata ancestry greater than 0.5, indicating preferential back-crossing to this parent.*

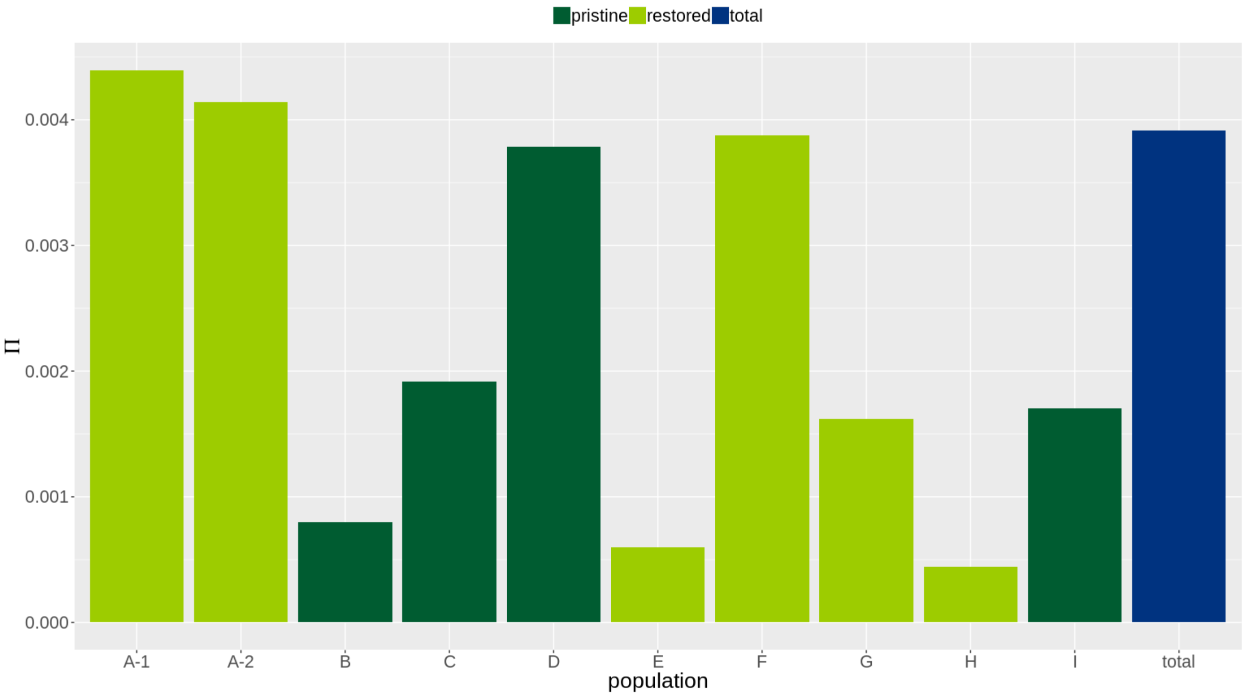


*Figure S 6: Barplot of average pairwise genetic diversity (π) within each site of both species combined including hybrids. Bar-color indicates the type of site.*

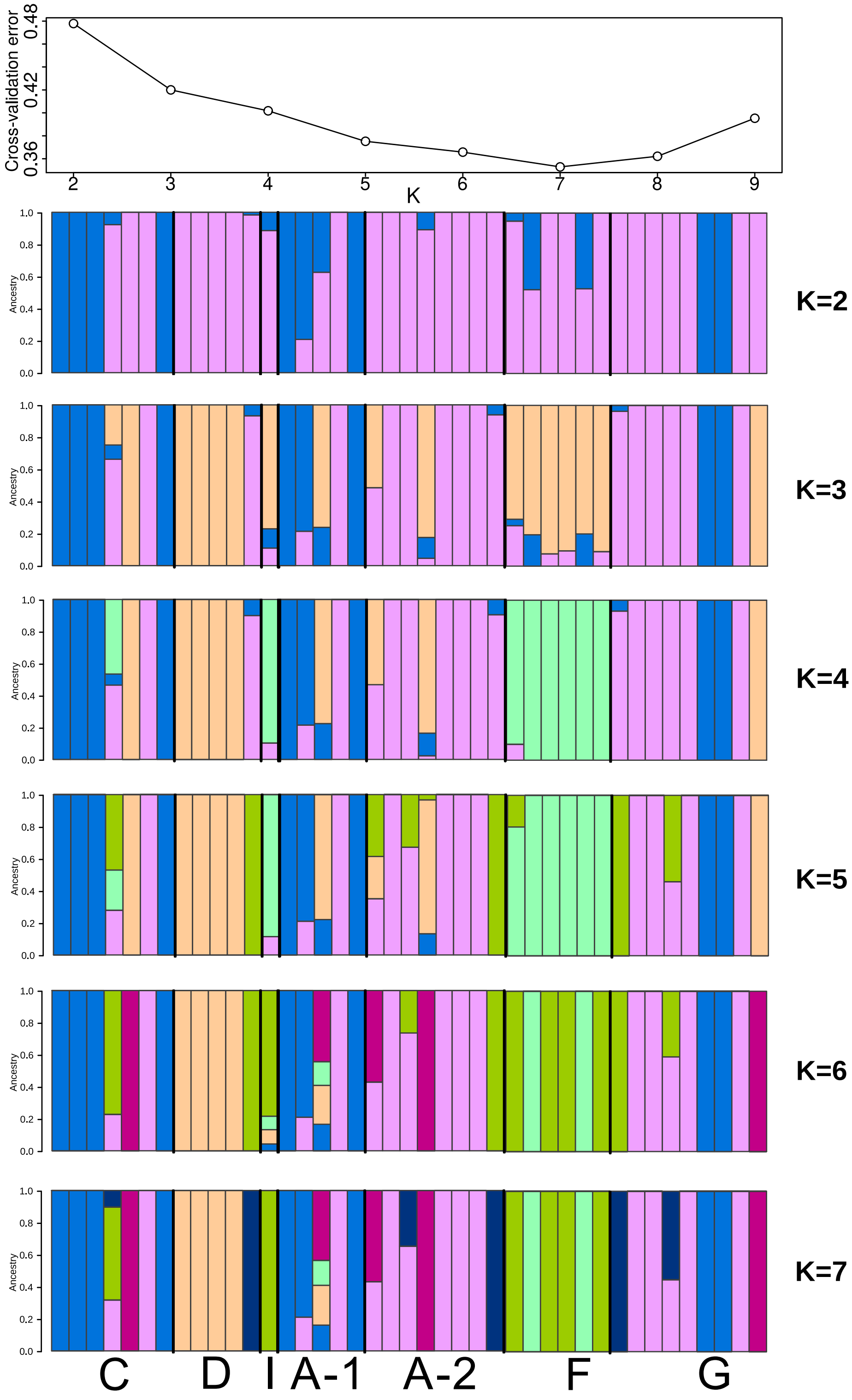


*Figure S 7: Arabis nemorensis ADMIXTURE analysis based on de novo genotypes for all K=2 until K=7.*

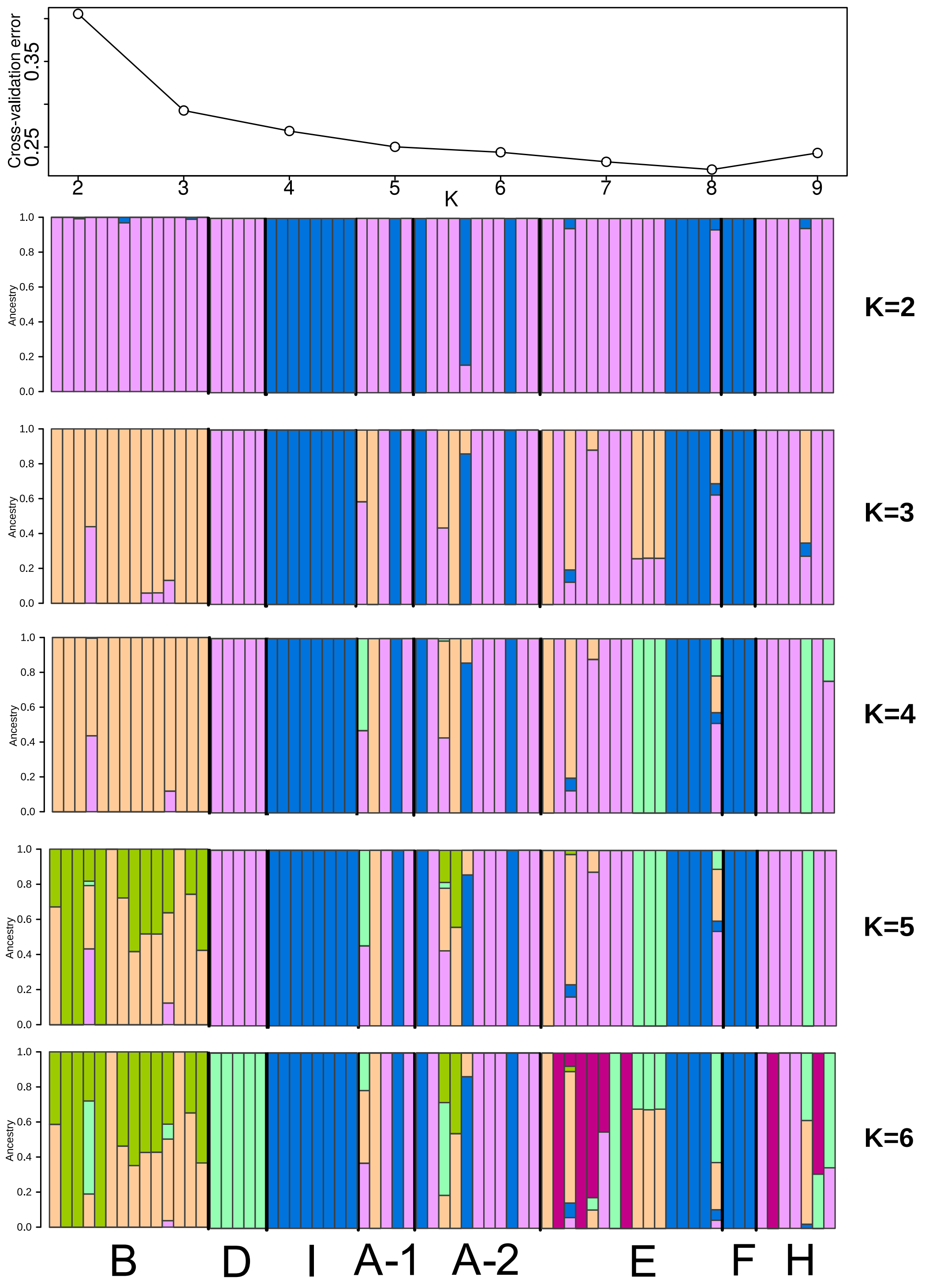


*Figure S 8: Arabis sagittata ADMIXTURE analysis based on de novo genotypes for all K=2 until K=6.*
