## Supplementary material for "Strengths and potential pitfalls of hay-transfer for ecological restoration revealed by RAD-seq analysis in floodplain *Arabis* species": Document S1

### Supplementary methods

#### Draft genome assembly and annotation

To facilitate genotype calling we assembled a draft genome for one *A. nemorensis* accession (ID 29). Library preparation and sequencing was done at the Cologne Center for Genomics. Details about the sequencing, assembly and annotation of the genome are described in Document S1. Three libraries were created: one paired-end library with approximately 280 bp insert size creating an overlap of 20 bp with 150 bp reads, and two mate-pair libraries with 3 kbp and 6 kbp inserts, respectively. The paired-end library was created using the Illumina TruSeq® Nano DNA Library Prep Kit, with 2µg of input DNA, without PCR. The mate-pair libraries were created using the Illumina Nextera® Mate Pair Library Prep Kit, with 4 µg of input DNA and 10 cycles of PCR for the 3 kbp library and 15 cycles of PCR for the 6 kbp library. All libraries were sequenced together as 150bp paired-end reads as part of an Illumina HiSeq 4000 lane for a total of 66 Gbp.

We used *FastQC* (Andrews, 2010) to quality-check the resulting reads. We filtered the resulting reads to remove reads shorter than 100bp and trimmed Illumina adapters using *Cutadapt* (Martin, 2011). We assembled reads using the *ALLPATHS-LG* assembler (Gnerre et al., 2011) with default settings, running it on the CHEOPS high-performance computing cluster of the University of Cologne. The resulting contig assembly had a size of 199 Mbp and an N50 of 47 kbp. The scaffold assembly had a size of 206 Mbp and an N50 102 kbp. To further scaffold the genome, we generated 2.9 Gbp of PacBio sequence data. Library preparation and sequencing was done at the Max-Planck-Institute for Plant Breeding Research (Cologne, Germany). We scaffolded the genome using *OPERA-LG* with default settings (Gao, Bertrand, Chia, & Nagarajan, 2016). This increased the N50 to 150kbp. To achieve chromosome-level assembly, we made use of the available reference genome of *Arabis alpina* (Jiao et al., 2017; Willing et al., 2015). To facilitate whole-genome alignment, repetitive regions in the genome were masked using RepeatMasker (Smit, Hubley, & Green, 2013). We used 'brassicaceae' as the search term ('-species' option) for the RepeatMasker repeat-database. We created the pseudo-chromosome assembly using the *CoGe* website (Lyons & Freeling, 2008). We aligned our draft genome with the *Arabis alpina* genome using *SynMap2* (Haug-Baltzell, Stephens, Davey, Scheidegger, & Lyons, 2017) and afterwards performed synthenic path assembly (Lyons, Freeling, Kustu, & Inwood, 2011) to assemble the chromosomes. The pseudo-chromosomes had a total size of 192 Mbp and were used for all following analyses.

The size of the genome was estimated by flow-cytometry, which was performed commercially at Plant Cytometry Services. The estimated genome size was 274 Mbp. Thus, the assembly size was 70% of the genome size.

##### Annotation

To detect and annotate Transposable elements (TE) specific to *A. nemorensis* (not in the previously used database) we used the softwares RepeatModeler (Smit & Hubley, 2008) and RepeatMasker (Smit et al., 2013). The consensus repeat library was first constructed by RepeatModeler and then used by RepeatMasker to search TEs. Protein-coding genes were annotated by integrating predictions of *ab initio* gene annotation tools and alignments of homologous proteins. Three different tools including Augustus v3.2.3 (Stanke & Waack, 2003), GlimmerHMM v3.0 (Majoros, Pertea, & Salzberg, 2004) and SNAP v2013 (Korf, 2004) were used to predict the initial gene models. Protein sequences from *A. thaliana*, *A. lyrata* and *A. alpina* (Arabidopsis Genome Initiative, 2000; Hu et al., 2011; Willing et al., 2015) were aligned to the assembly by the tool Exonerate v2.2.0 (Slater & Birney, 2005). Then, the *ab initio* predictions and protein alignment hits were further combined to build the consensus gene models by the tool EVidenceModeler (EVM) v2012 (Haas et al., 2008). Finally, TE related genes in these models were annotated by checking the TE annotation, blastp (Altschul, Gish, Miller, Myers, & Lipman, 1990) alignments with Plant TE related proteins and blastp alignments with *A. thaliana* proteins. If a gene's protein sequence had blastp alignment identity and coverage both larger than 50% with a TE related protein, or at least 30% of its exon regions overlapped with TEs but had no good blastp hit (identity >50% and coverage >50%) from *A. thaliana* protein sequences, this gene would be marked as a TE related gene.

### RAD-seq Protocol

##### Adapted from Etter et al. 2011

This protocol is more compact than the one by (Etter, Bassham, Hohenlohe, Johnson, & Cresko, 2011). We recommend to also take a look into the original protocol for additional information.

#### 1. Material

a. Restriction endonuclease digestion
    i. Restriction enzyme (e.g. KpnI; NEB)
    ii. Genomic DNA: 250ng per sample; min. 12.5 ng/µl
b. P1 Adapter ligation
    i. NEB Buffer 2
    ii. rATP(Promega): 100mM
    iii. P1 Adapter 450nM (see Appendix 1)
    iv. Concentrated T4 DNA Ligase (NEB): 2,000,000 U/ml
c. Purfication
    i. QIAquick or MinElute PCR Purification Kit (Qiagen)
    ii. Ampure XP Beads
d. End Repair
    i. Quick Blunting Kit (NEB)
e. 3`-dA overhang addition
    i. NEB Buffer 2
    ii. dATP (Fermentas): 10mM
    iii. Klenow Fragment (3´ to 5´ exo-, NEB): 5,000 U/ml.
f. P2 Adapter ligation
    i. NEB Buffer 2
    ii. rATP: 100mM
    iii. P2 Adapter 10µM (see Appendix 1)
    iv. Concentrated T4 DNA Ligase
g. RAD tag Amplification Enrichment
    i. Phusion High-Fidelity PCR Master Mix with HF Buffer
    ii. Primer Mix 10µM (see Appendix 1)

#### 2. Methods

##### 2.1 Restriction endonuclease digestion:

1. Digest 250 ng of DNA for each sample individually following manufacturer's instructions in 25 µl reaction volume
2. Heat inactivate the enzyme if possible. If not, reaction has to be cleaned using QIAquick column **only** if cut sites are recreated by adapter ligation (depends on barcode sequence)

##### 2.3 P1 Adapter Ligation

1. In this step each digestion sample is ligated with an individually barcoded P1 Adapter using the sticky overhang created by the restriction enzyme
2. To each digest add:

   0.5µl 10X NEB Buffer (the same as used for digestion)
   1.5µl Barcoded P1 Adapter 450 nM for KpnI
   0.3µl rATP (100mM)
   0.25µl T4 DNA Ligase (2,000,000 U/ml)
   2.45µl $H_2O$ (ad 30 µl)

***Note: Add adapter before the enzyme to avoid re-ligation of the genomic DNA***

3. Incubate 30 min @ room temperature

4. Heat-inactivate for 10 min @ 65 °C, let cool to RT afterwards

##### 2.4. Sample multiplexing

1. Combine barcoded samples at desired ratios to create pools that will later be multiplexed with the second adapter (see Appendix 2). For each pool, make 100µl Aliquots. Use one aliquot per pool to complete the protocol and freeze the rest as backup.

The following steps in the protocol are done for each of the pools.

##### 2.5 DNA shearing

1. For each pool one aliquot (100µl) is fragmented for 7 cycles with 30 s shearing followed by 30 s break
2. Check on tape station: average size should be 500-700bp (predominantly smaller than 1kb)

##### 2.6 Ampure XP cleanup and size selection

1. This step purifies the samples and removes fragments shorter than 200bp, which are mostly adapter dimers
2. Add to each sample 136µl AmpureXP beads and 22µl EB Buffer
3. Mix well by pipetting 10 times
4. Incubate 15 minutes
5. Put on magnet and incubate 10 minutes

6. Remove the liquid carefully without disturbing the beads
7. Wash twice with 200µl Ethanol (riddle tubes carefully)
8. Remove all ethanol and dry for 15 minutes until beads are matte
9. Remove from magnet
10. Add 20µl EB and re-suspend beads
11. Put on magnet and transfer liquid without beads into fresh tube

##### 2.7. End repair

1. To eluate from previous step add:
   2.5 µl 10x Blunting Buffer
   2.5 µl dNTP mix (1mM)
   1 µl Blunt Enzyme Mix
2. Incubate at RT for 30 minutes
3. Purify using Ampure beads or QIAquick column and elute in 43µl EB

##### 2.8. 3'-dA overhang addition

1. To eluate from previous step add:
   5 µl 10X NEB Buffer 2
   1µl dATP
   3µl Klenow-fragment
2. Incubate at 37°C for 30 min and slowly cool to ambient temperature
3. Heat inactivation for 5 min at 70°C and hold on 4°C

##### 2.9. P2 Adapter ligation

1. To inactivated reaction from previous step add:
   1 µl P2 Adapter (10µM)
   0.5 µl rATP (100mM)
   0.5 µl T4 DNA Ligase
2. Incubate at room temperature for 30 minutes
3. Purify using Ampure beads or QIAquick column and elute in 52µl EB

##### 2.10. RAD tag amplification

1. To determine library quality a test amplification should be done:

10.5 µl H2O
12.5 µl Phusion High-Fidelity Master Mix
1.0 µl Primer Mix (10µM); different mix for each pool as primer contains second barcode (index)
1.0 µl RAD library template (eluate from last step)
Run on Thermocycler with following program:
30 s @ 98° C
16x {10s @ 98° C; 30s @ 65° C; 30s @ 72° C}
5 min 72° C

Analyze the purified product on tapestation next to template to check if amplification worked.

2. If amplification worked, repeat the PCR under the same conditions but in 100µl volume with 4µl template, purify the product using Ampure Beads excluding fragments smaller than 200bp and check the results on tapestation. **Note: If amplification does not work well more template can be used, but more template also introduces more fragments that don't amplify, can't be sequenced and might disturb the sequencing process.**

3. Adjust the number of cycles if needed or try adding more template to the reaction.

4. Mix the indexed pools in desired ratios before sequencing.

#### Appendix

##### 1. Adapters

RADseq uses modified Illumina adapters with specific overhang for the restriction enzyme cut site. They also include barcodes (XXXXXX) for multiplexing and short random sequences (NNNNN) for identification of PCR duplicates. Each adapter is made from two custom oligos:

P1 with SacII-specific overhang:

5' - ACACTCTTTCCCTACACGACGCTCTTCCGATCTNNNNNXXXXXXG*C -3'

5' - [PHO]XXXXXXNNNNNAGATCGGAAGAGCGTCGTGTAGGGAAAGAGTGT - 3'

P2 universal:

5' - [PHO]AGATCGGAAGAGCGAGAACA*A -3'

5'- GTGACTGGAGTTCAGACGTGTGCTCTTCCGATCT*T -3'

Oligos should be HPLC purified. Best is NGSgrade oligos by Eurofins.

To anneal the complementary oligos, follow these instructions:

1. Prepare 100µM stocks for each oligo in 1X EB
2. Combine complementary oligos at 10µM in 1X Annealing Buffer (AB; 10X AB: 500mM NaCl, 100mM Tris-Cl, pH 7.5-8.0): 80µl AB, 10µl Oligo 1, 10µl Oligo 2
3. Run samples on Thermocycler with the following program: 2.5 min @ 97.5°C; cool to 21° C with -1.5 °C per minute; 1 min @ 21 °C; hold @ 4° C
4. Dilute to desired concentrations in AB and/or freeze as stock solution

**Final P1-adapter concentration and amount used in ligation depends on the number of DNA fragments (cut-sites).** I used the molarity calculator provided in the following publication: https://doi.org/10.1371/journal.pone.0037135

##### 2. Primers

RADseq uses modified Illumina primers. The reverse primer contains an index sequence that can be used for multiplexing. Indices are the same as used in Illumina kits.

Primer forward:

5’ – AATGATACGGCGACCACCGAGATCTACACTCTTTCCCTACACGACG -3’

Primer reverse:

5’ – CAAGCAGAAGACGGCATACGAGATXXXXXXGTGACTGGAGTTCAGACGTGTGC -3’

For the protocol prepare 10µM primer mixes with EB, e.g.: 80µl EB, 10µl Primer 1, 10µl Primer 2. Note that a specific primer mix with a different index is needed for each pool.

Primer sequences were taken from (Peterson, Weber, Kay, Fisher, & Hoekstra, 2012).

#### Complete list of primers from (Peterson et al., 2012):

| Name | OligoSequence |
|---|---|
| PCR1 | AATGATACGGCGACCACCGAGATCTACACTCTTTCCCTACACGACG |
| PCR2_Idx_1_ATCACG | CAAGCAGAAGACGGCATACGAGATCGTGATGTGACTGGAGTTCAGACGTGTGC |
| PCR2_Idx_2_CGATGT | CAAGCAGAAGACGGCATACGAGATACATCGGTGACTGGAGTTCAGACGTGTGC |
| PCR2_Idx_3_TTAGGC | CAAGCAGAAGACGGCATACGAGATGCCTAAGTGACTGGAGTTCAGACGTGTGC |
| PCR2_Idx_4_TGACCA | CAAGCAGAAGACGGCATACGAGATTGGTCAGTGACTGGAGTTCAGACGTGTGC |
| PCR2_Idx_5_ACAGTG | CAAGCAGAAGACGGCATACGAGATCACTGTGTGACTGGAGTTCAGACGTGTGC |
| PCR2_Idx_6_GCCAAT | CAAGCAGAAGACGGCATACGAGATATTGGCGTGACTGGAGTTCAGACGTGTGC |
| PCR2_Idx_7_CAGATC | CAAGCAGAAGACGGCATACGAGATGATCTGGTGACTGGAGTTCAGACGTGTGC |
| PCR2_Idx_8_ACTTGA | CAAGCAGAAGACGGCATACGAGATTCAAGTGTGACTGGAGTTCAGACGTGTGC |
| PCR2_Idx_9_GATCAG | CAAGCAGAAGACGGCATACGAGATCTGATCGTGACTGGAGTTCAGACGTGTGC |
| PCR2_Idx_10_TAGCTT | CAAGCAGAAGACGGCATACGAGATAAGCTAGTGACTGGAGTTCAGACGTGTGC |
| PCR2_Idx_11_GGCTAC | CAAGCAGAAGACGGCATACGAGATGTAGCCGTGACTGGAGTTCAGACGTGTGC |
| PCR2_Idx_12_CTTGTA | CAAGCAGAAGACGGCATACGAGATTACAAGGTGACTGGAGTTCAGACGTGTGC |
