## Supplementary material for "Strengths and potential pitfalls of hay-transfer for ecological restoration revealed by RAD-seq analysis in floodplain *Arabis* species": Document S2

Supplemental R Markdown Script


### Supplemental R Markdown Script

###### Hannes Dittberner

###### March 25, 2019

- 1 Introduction
- 2 Libraries and custom functions
- 3 Anaylsis of all samples
  - 3.1 Load reference based data
  - 3.2 Determine SNP types (coding/syn/nonsyn)
  - 3.3 Load stacks data
  - 3.4 Perform and plot PCA for both datasets
    - 3.4.1 Reference-based
    - 3.4.2 de-novo-based
  - 3.5 Delimit species and hybrids
  - 3.6 Diversity of both species combined
- 4 Analysis of species separately
  - 4.1 Load data
    - 4.1.1 Load reference based data for A. nemorensis
  - 4.2 Determine SNP types (coding/syn/nonsyn)
    - 4.2.1 Load stacks data for A. nemorensis
    - 4.2.2 Load reference data for A. sagittata
  - 4.3 Determine SNP types (coding/syn/nonsyn)
    - 4.3.1 Load stacks data for A. sagittata
  - 4.4 Calculation of pairwise Fst and genetic distance
    - 4.4.1 A. nemorensis reference-based
    - 4.4.2 A. sagittata reference-based
    - 4.4.3 Comparison of stacks- and reference-based estimates
  - 4.5 Calculation of genetic diversity within populations
    - 4.5.1 A. nemorensis reference-based
    - 4.5.2 A.sagittata reference-based
    - 4.5.3 A. nemorensis de-novo-based
    - 4.5.4 A. sagittata de-novo-based
    - 4.5.5 Comparison of reference-based and de-novo-based diversity estimates
  - 4.6 Admixture analysis
    - 4.6.1 A.nemorensis reference-based
    - 4.6.2 A. nemorensis de novo
    - 4.6.3 A. sagittata reference-based
    - 4.6.4 A. sagittata de novo
- 5 Map of populations

### 1 Introduction

This is an R-Markdown script describing the statistical analysis presented in the article. Figure names correspond to the ones used in the article and supplemental material. Additional figures were not named.

### 2 Libraries and custom functions

```
library(pegas)
library(hierfstat)
library(adegenet)
library(vcfR)
library(ggplot2)
library(ape)
library(heatmap3)
library(broom)
library(ggmap)
library(ggsn)
library(PopGenome)

rename_DNAbin = function(rows) {
    for (k in 1:length(rows)) {
        name = rows[k]
        name = unlist(strsplit(name, "_"))[1]
        name = sub("arabis", "", name)
        rows[k] = name
    }
    return(rows)
}
harmonic_number = function(n) {
    vec = numeric(n)
    for (j in 1:n) {
        vec[j] = 1/j
    }
    harm = sum(vec)
    return(harm)
}

diversity_statistics = function(vcf, GT_POS, unique_pops, pop_table) {
    
    dna = vcfR2DNAbin(vcf, unphased_as_NA = F, extract.haps = T)
    rownames(dna) = rename_DNAbin(rownames(dna))
    diversity = data.frame(population = rep(NA, length(unique_pops) + 1), pi = rep(NA, 
        length(unique_pops) + 1), theta = rep(NA, length(unique_pops) + 1), 
        D = rep(NA, length(unique_pops) + 1), D_pval = rep(NA, length(unique_pops) + 
            1), seg_sites = rep(NA, length(unique_pops) + 1), num_seqs = rep(NA, 
            length(unique_pops) + 1))
    sfss = list()
    for (i in 1:length(unique_pops)) {
        curr_pop = unique_pops[i]
        curr_indv = pop_table$V1[which(pop_table$V2 == curr_pop)]
        curr_dna = dna[which(rownames(dna) %in% curr_indv), ]
        if (nrow(curr_dna) == 0) {
            next
        }
        curr_div = (nuc.div(curr_dna, pairwise.deletion = T) * ncol(curr_dna))/GT_POS
        curr_theta = (length(seg.sites(curr_dna))/harmonic_number(nrow(curr_dna) - 
            1))/GT_POS
        curr_D = tajima.test(curr_dna)
        curr_seg_sites = seg.sites(curr_dna)
        diversity$population[i] = as.character(curr_pop)
        diversity$pi[i] = curr_div
        diversity$theta[i] = curr_theta
        diversity$D[i] = curr_D$D
        diversity$D_pval[i] = curr_D$Pval.beta
        diversity$seg_sites[i] = length(curr_seg_sites)
        diversity$num_seqs[i] = nrow(curr_dna)
        sfs = site.spectrum(curr_dna)
        class(sfs) = curr_pop
        sfss[[i]] = sfs
    }
    
    total_div = (nuc.div(dna, pairwise.deletion = T) * ncol(dna))/GT_POS
    total_theta = (length(seg.sites(dna))/harmonic_number(nrow(dna) - 1))/GT_POS
    total_D = tajima.test(dna)
    total_seg_sites = seg.sites(dna)
    diversity$population[i + 1] = "total"
    diversity$pi[i + 1] = total_div
    diversity$theta[i + 1] = total_theta
    diversity$D[i + 1] = total_D$D
    diversity$D_pval[i + 1] = total_D$Pval.beta
    diversity$seg_sites[i + 1] = length(total_seg_sites)
    sfs = site.spectrum(dna)
    class(sfs) = "total"
    sfss[[i]] = sfs
    
    outlist = list(divstats = diversity, sfs = sfss)
    return(outlist)
}

make_sidecolors = function(dist_mat, geodata) {
    rsidecolors = character(0)
    for (i in 1:nrow(dist_mat)) {
        pop = rownames(dist_mat)[i]
        status = geodata$status[geodata$Population == pop]
        if (status == "restored") {
            rsidecolors = c(rsidecolors, COLORS[8])
        } else if (status == "pristine") {
            rsidecolors = c(rsidecolors, COLORS[4])
        }
    }
    return(rsidecolors)
}

get_nonsyn_stats = function(VCF, GFF) {
    ns_stats = data.frame(CHROM = character(0), coding = numeric(0), syn = numeric(0), 
        nonsyn = numeric(0), total = numeric(0), stringsAsFactors = F, check.names = F)
    for (i in 1:8) {
        curr_CHROM = as.character(i)
        nemosag_class = readVCF(VCF, tid = curr_CHROM, frompos = 1, topos = 9e+08, 
            numcols = 10000, gffpath = GFF, include.unknown = T)
        nemosag_class = set.synnonsyn(nemosag_class, ref.chr = sub("LLLL", curr_CHROM, 
            "/home/hditt/Arabis/genome/pseudoassembly/pseudoassembly_chrLLLL.fa"))
        ns = nemosag_class@region.data@synonymous[[1]]
        stats = data.frame(CHROM = curr_CHROM, coding = nemosag_class@Coding.region, 
            syn = sum(ns == 1, na.rm = T), nonsyn = sum(ns == 0, na.rm = T), 
            total = nemosag_class@n.sites2, stringsAsFactors = F)
        ns_stats = rbind(ns_stats, stats)
    }
    ns_stats = rbind(ns_stats, data.frame(CHROM = "total", coding = sum(ns_stats$coding), 
        syn = sum(ns_stats$syn), nonsyn = sum(ns_stats$nonsyn), total = sum(ns_stats$total)))
    ns_stats = rbind(ns_stats, data.frame(CHROM = "total%", coding = sum(ns_stats$coding)/sum(ns_stats$total) * 
        100, syn = sum(ns_stats$syn)/sum(ns_stats$coding) * 100, nonsyn = sum(ns_stats$nonsyn)/sum(ns_stats$coding) * 
        100, total = NA))
    return(ns_stats)
}
```

### 3 Anaylsis of all samples

#### 3.1 Load reference based data

```
populations = read.table("/home/hditt/Arabis/samples_rad3/popmap_rhine.tsv", 
    sep = "\t", header = F, stringsAsFactors = F)
COLORS = c("#F0A3FF", "#0075DC", "#4C005C", "#005C31", "#2BCE48", "#FFCC99", 
    "#993F00", "#9DCC00", "#C20088", "#003380", "#FFA405", "#FFA8BB", "#00998F", 
    "#740AFF")
popdata = read.table("/home/hditt/Arabis/restoration_data/populations.csv", 
    sep = ",", header = T)

vcf_nemosag = read.vcfR("/home/hditt/Arabis/restoration_data/data/AllSamples_snps.vcf.gz")
```

```
## Scanning file to determine attributes.
## File attributes:
##   meta lines: 23
##   header_line: 24
##   variant count: 32880
##   column count: 143
## 
Meta line 23 read in.
## All meta lines processed.
## gt matrix initialized.
## Character matrix gt created.
##   Character matrix gt rows: 32880
##   Character matrix gt cols: 143
##   skip: 0
##   nrows: 32880
##   row_num: 0
## 
Processed variant 1000
Processed variant 2000
Processed variant 3000
Processed variant 4000
Processed variant 5000
Processed variant 6000
Processed variant 7000
Processed variant 8000
Processed variant 9000
Processed variant 10000
Processed variant 11000
Processed variant 12000
Processed variant 13000
Processed variant 14000
Processed variant 15000
Processed variant 16000
Processed variant 17000
Processed variant 18000
Processed variant 19000
Processed variant 20000
Processed variant 21000
Processed variant 22000
Processed variant 23000
Processed variant 24000
Processed variant 25000
Processed variant 26000
Processed variant 27000
Processed variant 28000
Processed variant 29000
Processed variant 30000
Processed variant 31000
Processed variant 32000
Processed variant: 32880
## All variants processed
```

```
vcf_nemosag@fix[, 3] = NA
loci_nemosag = vcfR2loci(vcf_nemosag)
genotypes_nemosag = t(extract.gt(vcf_nemosag))
colnames(genotypes_nemosag) = as.character(1:ncol(genotypes_nemosag))
variants_nemosag = df2genind(genotypes_nemosag, sep = "/")
populations_nemosag = populations[which(populations$V1 %in% row.names(variants_nemosag$tab)), 
    ]
pops_nemosag = populations_nemosag$V2[match(row.names(variants_nemosag$tab), 
    populations_nemosag$V1)]
variants_nemosag$pop = as.factor(pops_nemosag)
levels(variants_nemosag$pop) = c("A-1", "B", "A-2", "C", "D", "E", "F", "G", 
    "H", "I")
```

#### 3.2 Determine SNP types (coding/syn/nonsyn)

```
ns_stats_nemosag = get_nonsyn_stats("/home/hditt/Arabis/restoration_data/data/AllSamples_snps.vcf.gz", 
    "/home/hditt/Arabis/genome/pseudoassembly/Ane.protein-coding.genes.1.0.Add_ortholog.gff")
```

```
## 
## GFF information ...
## vcff::open : file opened, contains 134 samples
## [1] "Available ContigIdentifiers (parameter tid):"
## [1] "8" "1" "3" "4" "7" "5" "6" "2"
## |            :            |            :            | 100 %
## |====================================================
## 
## GFF information ...
## vcff::open : file opened, contains 134 samples
## [1] "Available ContigIdentifiers (parameter tid):"
## [1] "8" "1" "3" "4" "7" "5" "6" "2"
## |            :            |            :            | 100 %
## |====================================================
## 
## GFF information ...
## vcff::open : file opened, contains 134 samples
## [1] "Available ContigIdentifiers (parameter tid):"
## [1] "8" "1" "3" "4" "7" "5" "6" "2"
## |            :            |            :            | 100 %
## |====================================================
## 
## GFF information ...
## vcff::open : file opened, contains 134 samples
## [1] "Available ContigIdentifiers (parameter tid):"
## [1] "8" "1" "3" "4" "7" "5" "6" "2"
## |            :            |            :            | 100 %
## |====================================================
## 
## GFF information ...
## vcff::open : file opened, contains 134 samples
## [1] "Available ContigIdentifiers (parameter tid):"
## [1] "8" "1" "3" "4" "7" "5" "6" "2"
## |            :            |            :            | 100 %
## |====================================================
## 
## GFF information ...
## vcff::open : file opened, contains 134 samples
## [1] "Available ContigIdentifiers (parameter tid):"
## [1] "8" "1" "3" "4" "7" "5" "6" "2"
## |            :            |            :            | 100 %
## |====================================================
## 
## GFF information ...
## vcff::open : file opened, contains 134 samples
## [1] "Available ContigIdentifiers (parameter tid):"
## [1] "8" "1" "3" "4" "7" "5" "6" "2"
## |            :            |            :            | 100 %
## |====================================================
## 
## GFF information ...
## vcff::open : file opened, contains 134 samples
## [1] "Available ContigIdentifiers (parameter tid):"
## [1] "8" "1" "3" "4" "7" "5" "6" "2"
## |            :            |            :            | 100 %
## |====================================================
```

```
ns_stats_nemosag
```

| CHROM | coding | syn | nonsyn | total |
| --- | --- | --- | --- | --- |
| 1 | 1057.00000 | 436.00000 | 560.00000 | 4587 |
| 2 | 714.00000 | 267.00000 | 409.00000 | 3208 |
| 3 | 919.00000 | 307.00000 | 555.00000 | 4028 |
| 4 | 967.00000 | 374.00000 | 555.00000 | 4183 |
| 5 | 619.00000 | 229.00000 | 341.00000 | 3335 |
| 6 | 494.00000 | 179.00000 | 284.00000 | 3015 |
| 7 | 1022.00000 | 470.00000 | 530.00000 | 3980 |
| 8 | 1447.00000 | 644.00000 | 803.00000 | 6544 |
| total | 7239.00000 | 2906.00000 | 4037.00000 | 32880 |
| total% | 22.01642 | 40.14367 | 55.76737 | NA |

Table with SNP statistics for each chromosome and the whole genome. % Synonymous and non-synonymous are relative to the number of coding SNPs

#### 3.3 Load stacks data

```
stacks_all = read.vcfR("/home/hditt/Arabis/restoration_data/data/AllSamples_stacksSNPs.vcf")
```

```
## Scanning file to determine attributes.
## File attributes:
##   meta lines: 14
##   header_line: 15
##   variant count: 26467
##   column count: 143
## 
Meta line 14 read in.
## All meta lines processed.
## gt matrix initialized.
## Character matrix gt created.
##   Character matrix gt rows: 26467
##   Character matrix gt cols: 143
##   skip: 0
##   nrows: 26467
##   row_num: 0
## 
Processed variant 1000
Processed variant 2000
Processed variant 3000
Processed variant 4000
Processed variant 5000
Processed variant 6000
Processed variant 7000
Processed variant 8000
Processed variant 9000
Processed variant 10000
Processed variant 11000
Processed variant 12000
Processed variant 13000
Processed variant 14000
Processed variant 15000
Processed variant 16000
Processed variant 17000
Processed variant 18000
Processed variant 19000
Processed variant 20000
Processed variant 21000
Processed variant 22000
Processed variant 23000
Processed variant 24000
Processed variant 25000
Processed variant 26000
Processed variant: 26467
## All variants processed
```

```
stacks_all@fix[, 3] = NA
genotypes_stacks_all = t(extract.gt(stacks_all))
colnames(genotypes_stacks_all) = as.character(1:ncol(genotypes_stacks_all))
variants_stacks_all = df2genind(genotypes_stacks_all, sep = "/")
populations_rhine = populations[which(populations$V1 %in% row.names(variants_stacks_all$tab)), 
    ]
pops_rhine = populations_rhine$V2[match(row.names(variants_stacks_all$tab), 
    populations_rhine$V1)]
variants_stacks_all$pop = as.factor(pops_rhine)
levels(variants_stacks_all$pop) = c("A-1", "B", "A-2", "C", "D", "E", "F", "G", 
    "H", "I")
```

#### 3.4 Perform and plot PCA for both datasets

##### 3.4.1 Reference-based

```
variants_scaled_nemosag = scaleGen(variants_nemosag, NA.method = "mean")
pca_nemosag <- dudi.pca(variants_scaled_nemosag, cent = T, scale = F, scannf = F, 
    nf = 4)
axis_nemosag = pca_nemosag$eig * 100/sum(pca_nemosag$eig)
barplot(axis_nemosag[1:20], main = "PCA % explained", ylab = "% variance explained", 
    xlab = "Principal component", cex.lab = 1.4, names = 1:20, cex.axis = 1.4, 
    cex.names = 1.4)
```

Barplot of % variance explained by each principal component

```
col_nemosag = c("#990000", "#0075DC", "#993F00", "#4C005C", "#005C31", "#2BCE48", 
    "#FF5005", "#9DCC00", "#C20088", "#003380", "#FFA405", "#FFA8BB", "#00998F", 
    "#740AFF")
s.class(-pca_nemosag$li, pop(variants_nemosag), xax = 1, yax = 2, cstar = 0, 
    cpoint = 3, cellipse = 1, grid = T, clabel = 1.5, axesell = F, col = transp(col_nemosag))
```

Figure 2a: Plot of PCA of genetic variation among all samples. Samples are colored by population. Labels are in the centroid of each population.

##### 3.4.2 de-novo-based

```
variants_stacks_all_scaled = scaleGen(variants_stacks_all, NA.method = "mean")
pca_stacks_all <- dudi.pca(variants_stacks_all_scaled, cent = T, scale = F, 
    scannf = F, nf = 10)
axis_stacks_all = pca_stacks_all$eig * 100/sum(pca_stacks_all$eig)
barplot(axis_stacks_all[1:20], main = "PCA % explained")
```

Barplot of % variance explained by each principal component

```
col_stacks = c("#990000", "#0075DC", "#993F00", "#4C005C", "#005C31", "#2BCE48", 
    "#FF5005", "#9DCC00", "#C20088", "#003380", "#FFA405", "#FFA8BB", "#00998F", 
    "#740AFF")
s.class(pca_stacks_all$li, pop(variants_stacks_all), xax = 1, yax = 2, cstar = 0, 
    cpoint = 3, cellipse = 1, grid = T, clabel = 1.5, axesell = F, col = transp(col_stacks))
```

Figure 2b: Plot of PCA of genetic variation among all samples. Samples are colored by population. Labels are in the centroid of each population.

#### 3.5 Delimit species and hybrids

```
pca_tab = (-pca_nemosag$li)
pca_tab$pop = variants_nemosag$pop
pca_tab$spec = "NA"
pca_tab$spec[pca_tab$Axis1 > 150] = "A. sagittata"
pca_tab$spec[pca_tab$Axis1 < (-250)] = "A. nemorensis"
pca_tab$spec[pca_tab$Axis1 > (-250) & pca_tab$Axis1 < 150] = "hybrid"

spec_tab = as.data.frame(table(pca_tab$pop, pca_tab$spec))
names(spec_tab) = c("population", "species", "indvs")
spec_tab$population = factor(spec_tab$population, levels = c("A-1", "A-2", "B", 
    "C", "D", "E", "F", "G", "H", "I"))

spec_tab
```

| population | species | indvs |
| --- | --- | --- |
| A-1 | A. nemorensis | 5 |
| B | A. nemorensis | 0 |
| A-2 | A. nemorensis | 8 |
| C | A. nemorensis | 7 |
| D | A. nemorensis | 5 |
| E | A. nemorensis | 0 |
| F | A. nemorensis | 6 |
| G | A. nemorensis | 9 |
| H | A. nemorensis | 0 |
| I | A. nemorensis | 1 |
| A-1 | A. sagittata | 5 |
| B | A. sagittata | 14 |
| A-2 | A. sagittata | 11 |
| C | A. sagittata | 0 |
| D | A. sagittata | 5 |
| E | A. sagittata | 16 |
| F | A. sagittata | 3 |
| G | A. sagittata | 1 |
| H | A. sagittata | 7 |
| I | A. sagittata | 8 |
| A-1 | hybrid | 0 |
| B | hybrid | 3 |
| A-2 | hybrid | 3 |
| C | hybrid | 2 |
| D | hybrid | 13 |
| E | hybrid | 1 |
| F | hybrid | 0 |
| G | hybrid | 0 |
| H | hybrid | 1 |
| I | hybrid | 0 |

Overview of the number of samples from each species in each population

```
ggplot(data = spec_tab, aes(x = population, y = indvs, fill = species)) + geom_bar(stat = "identity") + 
    scale_fill_manual(values = c("tomato2", "royalblue3", "plum")) + theme_minimal() + 
    theme(text = element_text(size = 20), legend.position = "top", legend.title = element_blank()) + 
    ylab("Number of samples")
```

Figure 2c: Proportion of individuals of each species in each population

#### 3.6 Diversity of both species combined

```
populations_nemosag2 = populations_nemosag
populations_nemosag2$V2 = as.factor(populations_nemosag2$V2)
levels(populations_nemosag2$V2) = c("A-1", "B", "A-2", "C", "D", "E", "F", "G", 
    "H", "I")
divstats_nemosag = diversity_statistics(vcf_nemosag, GT_POS = 3565032, unique_pops = levels(variants_nemosag$pop), 
    pop_table = populations_nemosag2)  # genotyped postions (GT_POS) extracted from AllSamples_fullGT.vcf.gz
```

```
## 
Variant 3 processed
```

```
divstats_nemosag = divstats_nemosag$divstats
divstats_nemosag$type = c("restored", "pristine", "restored", "pristine", "pristine", 
    "restored", "restored", "restored", "restored", "pristine", "total")

ggplot(data = divstats_nemosag, aes(x = population, y = pi, fill = type)) + 
    geom_bar(stat = "identity") + theme(legend.position = "top", legend.title = element_blank(), 
    text = element_text(size = 20)) + ylab(expression(Pi)) + scale_fill_manual(values = COLORS[c(4, 
    8, 10)]) + xlab("population")
```

Within population genetic diversity of all individuals of both species combined, including hybrids

```
wilcox.test(divstats_nemosag$pi[divstats_nemosag$type == "restored"], divstats_nemosag$pi[divstats_nemosag$type == 
    "pristine"])
```

```
## 
##  Wilcoxon rank sum test
## 
## data:  divstats_nemosag$pi[divstats_nemosag$type == "restored"] and divstats_nemosag$pi[divstats_nemosag$type == "pristine"]
## W = 13, p-value = 0.9143
## alternative hypothesis: true location shift is not equal to 0
```

```
(mean(divstats_nemosag$pi[divstats_nemosag$type == "restored"]) - mean(divstats_nemosag$pi[divstats_nemosag$type == 
    "pristine"]))/mean(divstats_nemosag$pi[divstats_nemosag$type == "pristine"])
```

```
## [1] 0.2238595
```

### 4 Analysis of species separately

#### 4.1 Load data

##### 4.1.1 Load reference based data for A. nemorensis

```
vcf_nemo_rhine = read.vcfR("/home/hditt/Arabis/restoration_data/data/Anemorensis_snps.vcf.gz")
```

```
## Scanning file to determine attributes.
## File attributes:
##   meta lines: 23
##   header_line: 24
##   variant count: 2746
##   column count: 50
## 
Meta line 23 read in.
## All meta lines processed.
## gt matrix initialized.
## Character matrix gt created.
##   Character matrix gt rows: 2746
##   Character matrix gt cols: 50
##   skip: 0
##   nrows: 2746
##   row_num: 0
## 
Processed variant 1000
Processed variant 2000
Processed variant: 2746
## All variants processed
```

```
vcf_nemo_rhine@fix[, 3] = NA
loci_nemo_rhine = vcfR2loci(vcf_nemo_rhine)
genotypes = t(extract.gt(vcf_nemo_rhine))
colnames(genotypes) = as.character(1:ncol(genotypes))
variants_nemo_rhine = df2genind(genotypes, sep = "/")
populations_nemo_rhine = populations[which(populations$V1 %in% row.names(variants_nemo_rhine$tab)), 
    ]
populations_nemo_rhine[populations_nemo_rhine == "A-2"] = "A_res"
pops_nemo_rhine = populations_nemo_rhine$V2[match(row.names(variants_nemo_rhine$tab), 
    populations_nemo_rhine$V1)]
pops_nemo_rhine[pops_nemo_rhine == "A-2"] = "A_res"
variants_nemo_rhine$pop = as.factor(pops_nemo_rhine)
levels(variants_nemo_rhine$pop) = c("A-1", "A-2", "C", "D", "F", "G", "I")
```

#### 4.2 Determine SNP types (coding/syn/nonsyn)

```
ns_stats_nemo = get_nonsyn_stats("//home/hditt/Arabis/restoration_data/data/Anemorensis_snps.vcf.gz", 
    "/home/hditt/Arabis/genome/pseudoassembly/Ane.protein-coding.genes.1.0.Add_ortholog.gff")
```

```
## 
## GFF information ...
## vcff::open : file opened, contains 41 samples
## [1] "Available ContigIdentifiers (parameter tid):"
## [1] "8" "1" "3" "4" "7" "5" "6" "2"
## |            :            |            :            | 100 %
## |====================================================
## 
## GFF information ...
## vcff::open : file opened, contains 41 samples
## [1] "Available ContigIdentifiers (parameter tid):"
## [1] "8" "1" "3" "4" "7" "5" "6" "2"
## |            :            |            :            | 100 %
## |====================================================
## 
## GFF information ...
## vcff::open : file opened, contains 41 samples
## [1] "Available ContigIdentifiers (parameter tid):"
## [1] "8" "1" "3" "4" "7" "5" "6" "2"
## |            :            |            :            | 100 %
## |====================================================
## 
## GFF information ...
## vcff::open : file opened, contains 41 samples
## [1] "Available ContigIdentifiers (parameter tid):"
## [1] "8" "1" "3" "4" "7" "5" "6" "2"
## |            :            |            :            | 100 %
## |====================================================
## 
## GFF information ...
## vcff::open : file opened, contains 41 samples
## [1] "Available ContigIdentifiers (parameter tid):"
## [1] "8" "1" "3" "4" "7" "5" "6" "2"
## |            :            |            :            | 100 %
## |====================================================
## 
## GFF information ...
## vcff::open : file opened, contains 41 samples
## [1] "Available ContigIdentifiers (parameter tid):"
## [1] "8" "1" "3" "4" "7" "5" "6" "2"
## |            :            |            :            | 100 %
## |====================================================
## 
## GFF information ...
## vcff::open : file opened, contains 41 samples
## [1] "Available ContigIdentifiers (parameter tid):"
## [1] "8" "1" "3" "4" "7" "5" "6" "2"
## |            :            |            :            | 100 %
## |====================================================
## 
## GFF information ...
## vcff::open : file opened, contains 41 samples
## [1] "Available ContigIdentifiers (parameter tid):"
## [1] "8" "1" "3" "4" "7" "5" "6" "2"
## |            :            |            :            | 100 %
## |====================================================
```

```
ns_stats_nemo
```

| CHROM | coding | syn | nonsyn | total |
| --- | --- | --- | --- | --- |
| 1 | 32.0000 | 11.00000 | 20.00000 | 247 |
| 2 | 50.0000 | 9.00000 | 41.00000 | 334 |
| 3 | 70.0000 | 24.00000 | 43.00000 | 359 |
| 4 | 69.0000 | 18.00000 | 49.00000 | 300 |
| 5 | 25.0000 | 10.00000 | 13.00000 | 166 |
| 6 | 37.0000 | 14.00000 | 22.00000 | 411 |
| 7 | 161.0000 | 64.00000 | 95.00000 | 503 |
| 8 | 92.0000 | 33.00000 | 56.00000 | 426 |
| total | 536.0000 | 183.00000 | 339.00000 | 2746 |
| total% | 19.5193 | 34.14179 | 63.24627 | NA |

Table with SNP statistics for each chromosome and the whole genome. % Synonymous and non-synonymous are relative to the number of coding SNPs

##### 4.2.1 Load stacks data for A. nemorensis

```
vcf_nemo_stacks = read.vcfR("/home/hditt/Arabis/restoration_data/data/Anemorensis_stacksSNPs.vcf")
```

```
## Scanning file to determine attributes.
## File attributes:
##   meta lines: 14
##   header_line: 15
##   variant count: 14142
##   column count: 50
## 
Meta line 14 read in.
## All meta lines processed.
## gt matrix initialized.
## Character matrix gt created.
##   Character matrix gt rows: 14142
##   Character matrix gt cols: 50
##   skip: 0
##   nrows: 14142
##   row_num: 0
## 
Processed variant 1000
Processed variant 2000
Processed variant 3000
Processed variant 4000
Processed variant 5000
Processed variant 6000
Processed variant 7000
Processed variant 8000
Processed variant 9000
Processed variant 10000
Processed variant 11000
Processed variant 12000
Processed variant 13000
Processed variant 14000
Processed variant: 14142
## All variants processed
```

```
vcf_nemo_stacks@fix[, 3] = NA
genotypes = t(extract.gt(vcf_nemo_stacks))
colnames(genotypes) = as.character(1:ncol(genotypes))
variants_nemo_stacks = df2genind(genotypes, sep = "/")
populations_nemo_stacks = populations[which(populations$V1 %in% row.names(variants_nemo_stacks$tab)), 
    ]
populations_nemo_stacks[populations_nemo_stacks == "A-2"] = "A_res"
pops_nemo_stacks = populations_nemo_stacks$V2[match(row.names(variants_nemo_stacks$tab), 
    populations_nemo_stacks$V1)]
pops_nemo_stacks[pops_nemo_stacks == "A-2"] = "A_res"
variants_nemo_stacks$pop = as.factor(pops_nemo_stacks)
levels(variants_nemo_stacks$pop) = c("A-1", "A-2", "C", "D", "F", "G", "I")
```

##### 4.2.2 Load reference data for A. sagittata

```
vcf_sag_rhine = read.vcfR("/home/hditt/Arabis/restoration_data/data/Asagittata_snps.vcf.gz")
```

```
## Scanning file to determine attributes.
## File attributes:
##   meta lines: 23
##   header_line: 24
##   variant count: 6366
##   column count: 79
## 
Meta line 23 read in.
## All meta lines processed.
## gt matrix initialized.
## Character matrix gt created.
##   Character matrix gt rows: 6366
##   Character matrix gt cols: 79
##   skip: 0
##   nrows: 6366
##   row_num: 0
## 
Processed variant 1000
Processed variant 2000
Processed variant 3000
Processed variant 4000
Processed variant 5000
Processed variant 6000
Processed variant: 6366
## All variants processed
```

```
vcf_sag_rhine@fix[, 3] = NA
loci_sag_rhine = vcfR2loci(vcf_sag_rhine)
genotypes = t(extract.gt(vcf_sag_rhine))
colnames(genotypes) = as.character(1:ncol(genotypes))
variants_sag_rhine = df2genind(genotypes, sep = "/")
populations_sag_rhine = populations[which(populations$V1 %in% row.names(variants_sag_rhine$tab)), 
    ]
populations_sag_rhine[populations_sag_rhine == "Rvt"] = "I_pris"
pops_sag_rhine = populations_sag_rhine$V2[match(row.names(variants_sag_rhine$tab), 
    populations_sag_rhine$V1)]
pops_sag_rhine[pops_sag_rhine == "Rvt"] = "I_pris"
variants_sag_rhine$pop = as.factor(pops_sag_rhine)

levels(variants_sag_rhine$pop) = c("A-1", "B", "A-2", "D", "E", "F", "G", "H", 
    "I")
variants_sag_rhine$pop = factor(variants_sag_rhine$pop, levels = c("A-1", "A-2", 
    "B", "D", "E", "F", "G", "H", "I"))
```

#### 4.3 Determine SNP types (coding/syn/nonsyn)

```
ns_stats_sag = get_nonsyn_stats("/home/hditt/Arabis/restoration_data/data/Asagittata_snps.vcf.gz", 
    "/home/hditt/Arabis/genome/pseudoassembly/Ane.protein-coding.genes.1.0.Add_ortholog.gff")
```

```
## 
## GFF information ...
## vcff::open : file opened, contains 70 samples
## [1] "Available ContigIdentifiers (parameter tid):"
## [1] "8" "1" "3" "4" "7" "5" "6" "2"
## |            :            |            :            | 100 %
## |====================================================
## 
## GFF information ...
## vcff::open : file opened, contains 70 samples
## [1] "Available ContigIdentifiers (parameter tid):"
## [1] "8" "1" "3" "4" "7" "5" "6" "2"
## |            :            |            :            | 100 %
## |====================================================
## 
## GFF information ...
## vcff::open : file opened, contains 70 samples
## [1] "Available ContigIdentifiers (parameter tid):"
## [1] "8" "1" "3" "4" "7" "5" "6" "2"
## |            :            |            :            | 100 %
## |====================================================
## 
## GFF information ...
## vcff::open : file opened, contains 70 samples
## [1] "Available ContigIdentifiers (parameter tid):"
## [1] "8" "1" "3" "4" "7" "5" "6" "2"
## |            :            |            :            | 100 %
## |====================================================
## 
## GFF information ...
## vcff::open : file opened, contains 70 samples
## [1] "Available ContigIdentifiers (parameter tid):"
## [1] "8" "1" "3" "4" "7" "5" "6" "2"
## |            :            |            :            | 100 %
## |====================================================
## 
## GFF information ...
## vcff::open : file opened, contains 70 samples
## [1] "Available ContigIdentifiers (parameter tid):"
## [1] "8" "1" "3" "4" "7" "5" "6" "2"
## |            :            |            :            | 100 %
## |====================================================
## 
## GFF information ...
## vcff::open : file opened, contains 70 samples
## [1] "Available ContigIdentifiers (parameter tid):"
## [1] "8" "1" "3" "4" "7" "5" "6" "2"
## |            :            |            :            | 100 %
## |====================================================
## 
## GFF information ...
## vcff::open : file opened, contains 70 samples
## [1] "Available ContigIdentifiers (parameter tid):"
## [1] "8" "1" "3" "4" "7" "5" "6" "2"
## |            :            |            :            | 100 %
## |====================================================
```

```
ns_stats_sag
```

| CHROM | coding | syn | nonsyn | total |
| --- | --- | --- | --- | --- |
| 1 | 183.00000 | 61.00000 | 116.00000 | 892 |
| 2 | 95.00000 | 39.00000 | 51.00000 | 401 |
| 3 | 196.00000 | 66.00000 | 123.00000 | 1077 |
| 4 | 179.00000 | 62.00000 | 109.00000 | 641 |
| 5 | 169.00000 | 80.00000 | 79.00000 | 697 |
| 6 | 87.00000 | 36.00000 | 51.00000 | 571 |
| 7 | 387.00000 | 171.00000 | 201.00000 | 1068 |
| 8 | 221.00000 | 92.00000 | 129.00000 | 1019 |
| total | 1517.00000 | 607.00000 | 859.00000 | 6366 |
| total% | 23.82972 | 40.01318 | 56.62492 | NA |

Table with SNP statistics for each chromosome and the whole genome. % Synonymous and non-synonymous are relative to the number of coding SNPs

##### 4.3.1 Load stacks data for A. sagittata

```
vcf_sag_stacks = read.vcfR("/home/hditt/Arabis/restoration_data/data/Asagittata_stacksSNPs.vcf")
```

```
## Scanning file to determine attributes.
## File attributes:
##   meta lines: 14
##   header_line: 15
##   variant count: 13776
##   column count: 79
## 
Meta line 14 read in.
## All meta lines processed.
## gt matrix initialized.
## Character matrix gt created.
##   Character matrix gt rows: 13776
##   Character matrix gt cols: 79
##   skip: 0
##   nrows: 13776
##   row_num: 0
## 
Processed variant 1000
Processed variant 2000
Processed variant 3000
Processed variant 4000
Processed variant 5000
Processed variant 6000
Processed variant 7000
Processed variant 8000
Processed variant 9000
Processed variant 10000
Processed variant 11000
Processed variant 12000
Processed variant 13000
Processed variant: 13776
## All variants processed
```

```
vcf_sag_stacks@fix[, 3] = NA
# loci_nemo_rhine=vcfR2loci(vcf_nemo_rhine)
genotypes = t(extract.gt(vcf_sag_stacks))
colnames(genotypes) = as.character(1:ncol(genotypes))
variants_sag_stacks = df2genind(genotypes, sep = "/")
populations_sag_stacks = populations[which(populations$V1 %in% row.names(variants_sag_stacks$tab)), 
    ]
populations_sag_stacks[populations_sag_stacks == "Rvt"] = "I_pris"
pops_sag_stacks = populations_sag_stacks$V2[match(row.names(variants_sag_stacks$tab), 
    populations_sag_stacks$V1)]
pops_sag_stacks[pops_sag_stacks == "Rvt"] = "I_pris"
variants_sag_stacks$pop = as.factor(pops_sag_stacks)

# variants_sag_stacks=variants_sag_stacks[variants_sag_stacks$pop!='G_res']

levels(variants_sag_stacks$pop) = c("A-1", "B", "A-2", "D", "E", "F", "G", "H", 
    "I")
variants_sag_stacks$pop = factor(variants_sag_stacks$pop, levels = c("A-1", 
    "A-2", "B", "D", "E", "F", "G", "H", "I"))
```

#### 4.4 Calculation of pairwise Fst and genetic distance

##### 4.4.1 A. nemorensis reference-based

```
FST_nemo_rhine = as.matrix(genet.dist(variants_nemo_rhine, method = "Nei87"))
rownames(FST_nemo_rhine) = levels(variants_nemo_rhine$pop)
colnames(FST_nemo_rhine) = levels(variants_nemo_rhine$pop)
FST_nemo_rhine[FST_nemo_rhine < 0] = 0

Gdist_nemo_rhine = as.matrix(genet.dist(variants_nemo_rhine, method = "Dch"))
rownames(Gdist_nemo_rhine) = levels(variants_nemo_rhine$pop)
colnames(Gdist_nemo_rhine) = levels(variants_nemo_rhine$pop)

sidecolors_nemo_fst = make_sidecolors(FST_nemo_rhine, popdata)
heatmap3(as.matrix(FST_nemo_rhine), symm = T, cexRow = 2, cexCol = 2, margins = c(6, 
    6), ColSideLabs = "pop. status", RowSideLabs = "pop. status", RowAxisColors = 1, 
    ColAxisColors = 1, RowSideColors = sidecolors_nemo_fst, ColSideColors = sidecolors_nemo_fst, 
    col = colorRampPalette(c("black", "navy", "firebrick1"))(20))
```

Figure 4c: Heatmap of pairwise Fst among populations of A. nemorensis. Population labels and side-bars are colored by population type.

```
sidecolors_nemo_gdist = make_sidecolors(Gdist_nemo_rhine, popdata)
heatmap3(as.matrix(Gdist_nemo_rhine), symm = T, cexRow = 2, cexCol = 2, margins = c(6, 
    6), ColSideLabs = "pop. status", RowSideLabs = "pop. status", RowAxisColors = 1, 
    ColAxisColors = 1, RowSideColors = sidecolors_nemo_gdist, ColSideColors = sidecolors_nemo_gdist, 
    col = colorRampPalette(c("black", "navy", "firebrick1"))(20))
```

Figure 4a: Heatmap of pairwise genetic distance among populations of A. nemorensis. Population labels and side-bars are colored by population type.

```
# Comparison of genetic distance/Fst estimates among pristine and restored
# sites
Gdist_nemo_rhine_res = as.dist(Gdist_nemo_rhine[c(1, 2, 5, 6), c(1, 2, 5, 6)])
Gdist_nemo_rhine_pris = as.dist(Gdist_nemo_rhine[c(3, 4, 7), c(3, 4, 7)])

FST_nemo_rhine_res = as.dist(FST_nemo_rhine[c(1, 2, 5, 6), c(1, 2, 5, 6)])
FST_nemo_rhine_pris = as.dist(FST_nemo_rhine[c(3, 4, 7), c(3, 4, 7)])

wilcox.test(FST_nemo_rhine_pris, FST_nemo_rhine_res)
```

```
## 
##  Wilcoxon rank sum test
## 
## data:  FST_nemo_rhine_pris and FST_nemo_rhine_res
## W = 14, p-value = 0.2619
## alternative hypothesis: true location shift is not equal to 0
```

```
wilcox.test(Gdist_nemo_rhine_pris, Gdist_nemo_rhine_res)
```

```
## 
##  Wilcoxon rank sum test
## 
## data:  Gdist_nemo_rhine_pris and Gdist_nemo_rhine_res
## W = 17, p-value = 0.04762
## alternative hypothesis: true location shift is not equal to 0
```

##### 4.4.2 A. sagittata reference-based

```
FST_sag_rhine = as.matrix(genet.dist(variants_sag_rhine, method = "Nei87"))
rownames(FST_sag_rhine) = levels(variants_sag_rhine$pop)
colnames(FST_sag_rhine) = levels(variants_sag_rhine$pop)
FST_sag_rhine[FST_sag_rhine < 0] = 0

Gdist_sag_rhine = as.matrix(genet.dist(variants_sag_rhine, method = "Dch"))
rownames(Gdist_sag_rhine) = levels(variants_sag_rhine$pop)
colnames(Gdist_sag_rhine) = levels(variants_sag_rhine$pop)

sidecolors_sag_fst = make_sidecolors(FST_sag_rhine, popdata)
heatmap3(as.matrix(FST_sag_rhine), symm = T, cexRow = 2, cexCol = 2, margins = c(6, 
    6), ColSideLabs = "pop. status", RowSideLabs = "pop. status", RowAxisColors = 1, 
    ColAxisColors = 1, RowSideColors = sidecolors_sag_fst, ColSideColors = sidecolors_sag_fst, 
    col = colorRampPalette(c("black", "navy", "firebrick1"))(20))
```

Figure 4d: Heatmap of pairwise Fst among populations of A. nemorensis. Population labels and side-bars are colored by population type.

```
sidecolors_sag_gdist = make_sidecolors(Gdist_sag_rhine, popdata)
heatmap3(as.matrix(Gdist_sag_rhine), symm = T, cexRow = 2, cexCol = 2, ColSideLabs = "pop. status", 
    RowSideLabs = "pop. status", RowAxisColors = 1, ColAxisColors = 1, RowSideColors = sidecolors_sag_gdist, 
    ColSideColors = sidecolors_sag_gdist, margins = c(6, 6), col = colorRampPalette(c("black", 
        "navy", "firebrick1"))(20))
```

Figure 4b: Heatmap of pairwise genetic distance among populations of A. sagittata. Population labels and side-bars are colored by population type.

```
# Comparison of genetic distance/Fst estimates among pristine and restored
# sites
Gdist_sag_rhine_res = as.dist(Gdist_sag_rhine[c(1, 2, 5, 6, 7, 8), c(1, 2, 5, 
    6, 7, 8)])
Gdist_sag_rhine_pris = as.dist(Gdist_sag_rhine[c(3, 4, 9), c(3, 4, 9)])

FST_sag_rhine_res = as.dist(FST_sag_rhine[c(1, 2, 5, 6, 7, 8), c(1, 2, 5, 6, 
    7, 8)])
FST_sag_rhine_pris = as.dist(FST_sag_rhine[c(3, 4, 9), c(3, 4, 9)])

wilcox.test(FST_sag_rhine_pris, FST_sag_rhine_res)
```

```
## 
##  Wilcoxon rank sum test with continuity correction
## 
## data:  FST_sag_rhine_pris and FST_sag_rhine_res
## W = 43, p-value = 0.01581
## alternative hypothesis: true location shift is not equal to 0
```

```
wilcox.test(Gdist_sag_rhine_pris, Gdist_sag_rhine_res)
```

```
## 
##  Wilcoxon rank sum test
## 
## data:  Gdist_sag_rhine_pris and Gdist_sag_rhine_res
## W = 45, p-value = 0.002451
## alternative hypothesis: true location shift is not equal to 0
```

##### 4.4.3 Comparison of stacks- and reference-based estimates

```
FST_nemo_stacks = as.matrix(genet.dist(variants_nemo_stacks, method = "Nei87"))
rownames(FST_nemo_stacks) = levels(variants_nemo_stacks$pop)
colnames(FST_nemo_stacks) = levels(variants_nemo_stacks$pop)
FST_nemo_stacks[FST_nemo_stacks < 0] = 0

Gdist_nemo_stacks = as.matrix(genet.dist(variants_nemo_stacks, method = "Dch"))
rownames(Gdist_nemo_stacks) = levels(variants_nemo_stacks$pop)
colnames(Gdist_nemo_stacks) = levels(variants_nemo_stacks$pop)

FST_nemo_table = tidy(as.dist(FST_nemo_rhine))
FST_nemo_stacks_table = tidy(as.dist(FST_nemo_stacks))
FST_nemo_table = cbind(FST_nemo_table, FST_nemo_stacks_table$distance, "A. nemorensis")
names(FST_nemo_table) = c("pop1", "pop2", "FST_ref", "FST_stacks", "species")

Gdist_nemo_table = tidy(as.dist(Gdist_nemo_rhine))
Gdist_nemo_stacks_table = tidy(as.dist(Gdist_nemo_stacks))

Gdist_nemo_table = cbind(Gdist_nemo_table, Gdist_nemo_stacks_table$distance, 
    "A. nemorensis")
names(Gdist_nemo_table) = c("pop1", "pop2", "Gdist_ref", "Gdist_stacks", "species")

FST_sag_stacks = as.matrix(genet.dist(variants_sag_stacks, method = "Nei87"))
rownames(FST_sag_stacks) = levels(variants_sag_stacks$pop)
colnames(FST_sag_stacks) = levels(variants_sag_stacks$pop)
FST_sag_stacks[FST_sag_stacks < 0] = 0

Gdist_sag_stacks = as.matrix(genet.dist(variants_sag_stacks, method = "Dch"))
rownames(Gdist_sag_stacks) = levels(variants_sag_stacks$pop)
colnames(Gdist_sag_stacks) = levels(variants_sag_stacks$pop)

FST_sag_table = tidy(as.dist(FST_sag_rhine))
FST_sag_stacks_table = tidy(as.dist(FST_sag_stacks))

FST_sag_table = cbind(FST_sag_table, FST_sag_stacks_table$distance, "A. sagittata")
names(FST_sag_table) = c("pop1", "pop2", "FST_ref", "FST_stacks", "species")

FST_all_table = rbind(FST_nemo_table, FST_sag_table)

Gdist_sag_table = tidy(as.dist(Gdist_sag_rhine))
Gdist_sag_stacks_table = tidy(as.dist(Gdist_sag_stacks))

Gdist_sag_table = cbind(Gdist_sag_table, Gdist_sag_stacks_table$distance, "A. sagittata")
names(Gdist_sag_table) = c("pop1", "pop2", "Gdist_ref", "Gdist_stacks", "species")

Gdist_all_table = rbind(Gdist_nemo_table, Gdist_sag_table)

# Mantel test of Fst estimates for A. nemorensis
mantel.randtest(as.dist(FST_nemo_stacks), as.dist(FST_nemo_rhine), nrepet = 10000)
```

```
## Monte-Carlo test
## Call: mantel.randtest(m1 = as.dist(FST_nemo_stacks), m2 = as.dist(FST_nemo_rhine), 
##     nrepet = 10000)
## 
## Observation: 0.9898102 
## 
## Based on 10000 replicates
## Simulated p-value: 0.00069993 
## Alternative hypothesis: greater 
## 
##      Std.Obs  Expectation     Variance 
## 3.4710694590 0.0006801332 0.0812045115
```

```
# Mantel test of Fst estimates for A. sagittata
mantel.randtest(as.dist(FST_sag_stacks), as.dist(FST_sag_rhine), nrepet = 1e+05)
```

```
## Monte-Carlo test
## Call: mantel.randtest(m1 = as.dist(FST_sag_stacks), m2 = as.dist(FST_sag_rhine), 
##     nrepet = 1e+05)
## 
## Observation: 0.9900522 
## 
## Based on 100000 replicates
## Simulated p-value: 1.99998e-05 
## Alternative hypothesis: greater 
## 
##     Std.Obs Expectation    Variance 
##  3.94038320  0.00155425  0.06293245
```

```
ggplot(data = FST_all_table, aes(x = FST_ref, y = FST_stacks, col = species)) + 
    geom_point(size = 4) + geom_smooth(method = "lm", fullrange = T) + theme_minimal() + 
    theme(legend.position = "top", legend.title = element_blank(), text = element_text(size = 20)) + 
    xlab("Fst reference") + ylab("Fst de novo") + scale_color_manual(values = c("tomato2", 
    "royalblue3")) + geom_abline(slope = 1, intercept = 0, lty = "dashed", size = 1.2)
```

Figure 5: Correlation of Fst estimates from the de novo and reference pipeline for each species. Lines are linear fit through the points and grey shades are the error of the fit.

```
# Mantel test of genetic distance estimates for A. nemorensis
mantel.randtest(as.dist(Gdist_nemo_stacks), as.dist(Gdist_nemo_rhine), nrepet = 10000)
```

```
## Monte-Carlo test
## Call: mantel.randtest(m1 = as.dist(Gdist_nemo_stacks), m2 = as.dist(Gdist_nemo_rhine), 
##     nrepet = 10000)
## 
## Observation: 0.997798 
## 
## Based on 10000 replicates
## Simulated p-value: 9.999e-05 
## Alternative hypothesis: greater 
## 
##     Std.Obs Expectation    Variance 
## 3.075450999 0.001816905 0.104878032
```

```
# Mantel test of genetic distance estimates for A. sagittata
mantel.randtest(as.dist(Gdist_sag_stacks), as.dist(Gdist_sag_rhine), nrepet = 1e+05)
```

```
## Monte-Carlo test
## Call: mantel.randtest(m1 = as.dist(Gdist_sag_stacks), m2 = as.dist(Gdist_sag_rhine), 
##     nrepet = 1e+05)
## 
## Observation: 0.9871029 
## 
## Based on 100000 replicates
## Simulated p-value: 9.9999e-06 
## Alternative hypothesis: greater 
## 
##      Std.Obs  Expectation     Variance 
## 4.4751442721 0.0006168126 0.0485923425
```

```
ggplot(data = Gdist_all_table, aes(x = Gdist_ref, y = Gdist_stacks, col = species)) + 
    geom_point(size = 4) + geom_smooth(method = "lm", fullrange = T) + theme_minimal() + 
    theme(legend.position = "top", legend.title = element_blank(), text = element_text(size = 20)) + 
    xlab("Genetic distance reference") + ylab("Genetic distance de novo") + 
    scale_color_manual(values = c("tomato2", "royalblue3")) + geom_abline(slope = 1, 
    intercept = 0, lty = "dashed", size = 1.2)
```

Figure 5: Correlation of genetic distance estimates from the de novo and reference pipeline for each species. Lines are linear fit through the points and grey shades are the error of the fit.

#### 4.5 Calculation of genetic diversity within populations

##### 4.5.1 A. nemorensis reference-based

```
diversity_nemo_rhine = diversity_statistics(vcf_nemo_rhine, 4704842, unique(pops_nemo_rhine), 
    populations_nemo_rhine)  # number of genotype positions (GT_pos) extracted from Anemorensis_fullGT.vcf.gz
```

```
## 
Variant 3 processed
```

```
diversity_nemo_rhine_sfs = diversity_nemo_rhine$sfs
diversity_nemo_rhine = diversity_nemo_rhine$divstats

diversity_nemo_rhine = diversity_nemo_rhine[diversity_nemo_rhine$population != 
    "Rvt", ]  #remove population with one individual
diversity_nemo_rhine = diversity_nemo_rhine[order(diversity_nemo_rhine$population), 
    ]
diversity_nemo_rhine$type = c("restored", "restored", "pristine", "pristine", 
    "restored", "restored", "total")
diversity_nemo_rhine$populationplot = as.factor(diversity_nemo_rhine$population)
levels(diversity_nemo_rhine$populationplot) = c("A-1", "A-2", "C", "D", "F", 
    "G", "total")
diversity_nemo_rhine$populationplot = factor(diversity_nemo_rhine$populationplot, 
    levels = c("C", "D", "A-1", "A-2", "F", "G", "total"))

ggplot(data = diversity_nemo_rhine, aes(x = populationplot, y = pi, fill = type)) + 
    geom_bar(stat = "identity") + theme(legend.position = "top", legend.title = element_blank(), 
    text = element_text(size = 20)) + ylab(expression(Pi)) + ylim(0, 4e-04) + 
    scale_fill_manual(values = COLORS[c(4, 8, 10)]) + xlab("population")
```

Figure 3a: Within population genetic diversity of A. nemorensis. Populations are colored by type.

```
# Test for difference in genetic diversity between pristine and restored
# sites
wilcox.test(diversity_nemo_rhine$pi[diversity_nemo_rhine$type == "restored"], 
    diversity_nemo_rhine$pi[diversity_nemo_rhine$type == "pristine"])
```

```
## 
##  Wilcoxon rank sum test
## 
## data:  diversity_nemo_rhine$pi[diversity_nemo_rhine$type == "restored"] and diversity_nemo_rhine$pi[diversity_nemo_rhine$type == "pristine"]
## W = 4, p-value = 1
## alternative hypothesis: true location shift is not equal to 0
```

##### 4.5.2 A.sagittata reference-based

```
diversity_sag_rhine = diversity_statistics(vcf_sag_rhine, 3619235, unique(pops_sag_rhine), 
    populations_sag_rhine)  # number of genotype positions (GT_pos) extracted from Asagittata_fullGT.vcf.gz
```

```
## 
Variant 3 processed
```

```
diversity_sag_rhine_sfs = diversity_sag_rhine$sfs
diversity_sag_rhine = diversity_sag_rhine$divstats

diversity_sag_rhine = diversity_sag_rhine[diversity_sag_rhine$population != 
    "G_res", ]  #remove population with one individual
diversity_sag_rhine = diversity_sag_rhine[order(diversity_sag_rhine$population), 
    ]
diversity_sag_rhine$type = c("restored", "pristine", "restored", "pristine", 
    "restored", "restored", "restored", "pristine", "total")

diversity_sag_rhine$populationplot = as.factor(diversity_sag_rhine$population)
levels(diversity_sag_rhine$populationplot) = c("A-1", "B", "A-2", "D", "E", 
    "F", "H", "I", "total")
diversity_sag_rhine$populationplot = factor(diversity_sag_rhine$populationplot, 
    levels = c("A-1", "A-2", "B", "D", "E", "F", "H", "I", "total"))

diversity_sag_rhine$populationplot = factor(diversity_sag_rhine$populationplot, 
    levels = c("B", "D", "I", "A-1", "A-2", "E", "F", "H", "total"))


ggplot(data = diversity_sag_rhine, aes(x = populationplot, y = pi, fill = type)) + 
    geom_bar(stat = "identity") + theme(legend.position = "top", legend.title = element_blank(), 
    text = element_text(size = 20)) + ylab(expression(Pi)) + scale_fill_manual(values = COLORS[c(4, 
    8, 10)]) + xlab("population")
```

Figure 3b: Within population genetic diversity of A. nemorensis. Populations are colored by type.

```
# Test for difference in genetic diversity between pristine and restored
# sites
wilcox.test(diversity_sag_rhine$pi[diversity_sag_rhine$type == "restored"], 
    diversity_sag_rhine$pi[diversity_sag_rhine$type == "pristine"])
```

```
## 
##  Wilcoxon rank sum test
## 
## data:  diversity_sag_rhine$pi[diversity_sag_rhine$type == "restored"] and diversity_sag_rhine$pi[diversity_sag_rhine$type == "pristine"]
## W = 13, p-value = 0.1429
## alternative hypothesis: true location shift is not equal to 0
```

##### 4.5.3 A. nemorensis de-novo-based

```
GT_POS_nemo_stacks = 15684223 - 66931  # total number of genotyped sites from populations.log
diversity_nemo_stacks = diversity_statistics(vcf_nemo_stacks, GT_POS_nemo_stacks, 
    unique(pops_nemo_stacks), pop_table = populations_nemo_stacks)
```

```
## 
Variant 3 processed
```

```
diversity_nemo_stacks_sfs = diversity_nemo_stacks$sfs
diversity_nemo_stacks = diversity_nemo_stacks$divstats
diversity_nemo_stacks = diversity_nemo_stacks[diversity_nemo_stacks$population != 
    "Rvt", ]  #remove population with one individual
```

##### 4.5.4 A. sagittata de-novo-based

```
GT_POS_sag_stacks = 13774559 - 68511  # total number of genotyped sites from populations.log
diversity_sag_stacks = diversity_statistics(vcf_sag_stacks, GT_POS_sag_stacks, 
    unique(pops_sag_stacks), populations_sag_stacks)
```

```
## 
Variant 3 processed
```

```
diversity_sag_stacks_sfs = diversity_sag_stacks$sfs
diversity_sag_stacks = diversity_sag_stacks$divstats

diversity_sag_stacks = diversity_sag_stacks[diversity_sag_stacks$population != 
    "G_res", ]  #remove population with one individual
```

##### 4.5.5 Comparison of reference-based and de-novo-based diversity estimates

```
diversity_nemo_rhine = diversity_nemo_rhine[order(diversity_nemo_rhine$population), 
    ]
diversity_all = data.frame(Population = diversity_nemo_rhine$population, Pi_ref = diversity_nemo_rhine$pi, 
    Pi_stacks = diversity_nemo_stacks$pi, Species = "A.nemorensis")

diversity_sag_rhine = diversity_sag_rhine[order(diversity_sag_rhine$population), 
    ]
diversity_all = rbind(diversity_all, data.frame(Population = diversity_sag_rhine$population, 
    Pi_ref = diversity_sag_rhine$pi, Pi_stacks = diversity_sag_stacks$pi, Species = "A. sagittata"))

ggplot(data = diversity_all, aes(x = Pi_ref, y = Pi_stacks, col = Species)) + 
    geom_point(size = 4) + geom_smooth(method = "lm", fullrange = T) + theme_minimal() + 
    theme(legend.position = "top", legend.title = element_blank(), text = element_text(size = 20)) + 
    scale_color_manual(values = c("tomato2", "royalblue3")) + xlab(expression(Pi ~ 
    "reference")) + ylab(expression(Pi ~ "denovo")) + geom_abline(slope = 1, 
    intercept = 0, lty = "dashed", size = 1.2)
```

Figure 5: Correlation of genetic diversity estimates from the de novo and reference pipeline for each species. Lines are linear fit through the points and grey shades are the error of the fit.

```
# Pearson Correlation test for genetic diversity estimates of two pipelines
# for A. nemorensis
cor.test(diversity_all$Pi_ref[diversity_all$Species == "A.nemorensis"], diversity_all$Pi_stacks[diversity_all$Species == 
    "A.nemorensis"])
```

```
## 
##  Pearson's product-moment correlation
## 
## data:  diversity_all$Pi_ref[diversity_all$Species == "A.nemorensis"] and diversity_all$Pi_stacks[diversity_all$Species == "A.nemorensis"]
## t = 7.0796, df = 5, p-value = 0.0008702
## alternative hypothesis: true correlation is not equal to 0
## 95 percent confidence interval:
##  0.7112519 0.9933261
## sample estimates:
##       cor 
## 0.9535665
```

```
# Pearson Correlation test for genetic diversity estimates of two pipelines
# for A. sagittata
cor.test(diversity_all$Pi_ref[diversity_all$Species == "A. sagittata"], diversity_all$Pi_stacks[diversity_all$Species == 
    "A. sagittata"])
```

```
## 
##  Pearson's product-moment correlation
## 
## data:  diversity_all$Pi_ref[diversity_all$Species == "A. sagittata"] and diversity_all$Pi_stacks[diversity_all$Species == "A. sagittata"]
## t = 43.195, df = 7, p-value = 9.305e-10
## alternative hypothesis: true correlation is not equal to 0
## 95 percent confidence interval:
##  0.9907661 0.9996222
## sample estimates:
##       cor 
## 0.9981294
```

#### 4.6 Admixture analysis

```
barNaming <- function(vec) {
    retVec <- as.character(vec)
    for (k in 2:length(vec)) {
        if (vec[k - 1] == vec[k]) 
            retVec[k] <- NA
    }
    retVecMid = rep(NA, length(retVec))
    starts = which(!is.na(retVec))
    for (i in 1:length(starts) - 1) {
        mid = starts[i] + floor((starts[i + 1] - starts[i] + 1)/2)
        retVecMid[mid] = as.character(retVec[starts[i]])
    }
    mid = starts[i + 1] + floor((length(retVec) - starts[i + 1] + 1)/2)
    retVecMid[mid] = as.character(retVec[starts[i + 1]])
    retVecMid = c(retVecMid[2:length(retVecMid)], NA)
    return(as.factor(retVecMid))
}

vertLinePos = function(vec) {
    retVec <- as.character(vec)
    for (k in 2:length(vec)) {
        if (vec[k - 1] == vec[k]) 
            retVec[k] <- NA
    }
    l = which(!is.na(retVec))
    return(l)
}
```

##### 4.6.1 A.nemorensis reference-based

```
fam_nemo_rhine = read.table("/home/hditt/Arabis/samples_rad3/variants/ArabisNemoRhine_pseudo_0.95miss_reprem_loci_hetrem20_snps.recode.vcf.fam")
populations_nemo = read.table("/home/hditt/Arabis/samples_rad3/variants/populations_nemo.txt", 
    header = T)
populations_nemo_rhine = populations_nemo[populations_nemo$V2 == "A_res" | populations_nemo$V2 == 
    "B_res" | populations_nemo$V2 == "B_pris" | populations_nemo$V2 == "C_pris" | 
    populations_nemo$V2 == "D_pris" | populations_nemo$V2 == "F_res" | populations_nemo$V2 == 
    "G_res" | populations_nemo$V2 == "Rvt", ]
populations_nemo_rhine$V2 = as.character(populations_nemo_rhine$V2)
populations_nemo_rhine$V2 = as.factor(populations_nemo_rhine$V2)
levels(populations_nemo_rhine$V2) = c("A-1", "A-2", "C", "D", "F", "G", "I")
populations_nemo_rhine$V2 = as.character(populations_nemo_rhine$V2)

cvtable = read.table("/home/hditt/Arabis/samples_rad3/variants/admix_NemoRhine_30052018/ArabisNemoRhine_pseudo_0.95miss_reprem_loci_hetrem20_snps.recode.vcf_CV.log", 
    sep = ",", header = T)
cvtable = cvtable[order(cvtable$K), ]

plot(cvtable$K, cvtable$CV, type = "b", xlab = "K", ylab = "Cross-validation error")
```

```
COLORS_mx = c("#FFF176", "#FFCA28", "#FF9800", "#E53935", "#C2185B", "#6A1B9A", 
    "#1A237E")
for (i in 2:7) {
    admix_table = read.table(sub("LLLL", as.character(i), "/home/hditt/Arabis/samples_rad3/variants/admix_NemoRhine_30052018/ArabisNemoRhine_pseudo_0.95miss_reprem_loci_hetrem20_snps.recode.vcf.LLLL.Q.converted"))
    admix_table = admix_table[match(populations_nemo_rhine$V1, fam_nemo_rhine$V1), 
        ]
    b = barplot(t(as.matrix(admix_table)), col = COLORS_mx, main = paste("K=", 
        as.character(i), sep = ""), ylab = "Ancestry", border = "grey35", space = 0, 
        names.arg = barNaming(populations_nemo_rhine$V2), las = 2, cex.names = 1.5, 
        cex.axis = 1.5, cex.lab = 1.5, cex.main = 2, horiz = F)
    abline(v = b[vertLinePos(populations_nemo_rhine$V2)] - 0.5, lwd = 8, col = "black")
}
```

Overview of ADMIXTURE plots from K=2 to K=7 for A. nemorensis based on the reference pipeline.

##### 4.6.2 A. nemorensis de novo

```
fam_nemo_rhine = read.table("/home/hditt/Arabis/samples_rad3/stacks/denovo_M6_n6_new/popstats_nemo/populations.snps.vcf.fam")

cvtable = read.table("/home/hditt/Arabis/samples_rad3/stacks/denovo_M6_n6_new/popstats_nemo/admix/populations.snps.vcf_CV.log", 
    sep = ",", header = T)
cvtable = cvtable[order(cvtable$K), ]

plot(cvtable$K, cvtable$CV, type = "b", xlab = "K", ylab = "Cross-validation error")
```

```
COLORS_mx = c("#F0A3FF", "#0075DC", "#FFCC99", "#94FFB5", "#9DCC00", "#C20088", 
    "#003380", "#FFA405", "#FFA8BB")
for (i in 2:7) {
    admix_table = read.table(sub("LLLL", as.character(i), "/home/hditt/Arabis/samples_rad3/stacks/denovo_M6_n6_new/popstats_nemo/admix/populations.snps.vcf.LLLL.Q.converted"))
    admix_table = admix_table[match(populations_nemo_rhine$V1, fam_nemo_rhine$V1), 
        ]
    b = barplot(t(as.matrix(admix_table)), col = COLORS_mx, main = paste("K=", 
        as.character(i), sep = ""), ylab = "Ancestry", border = "grey25", space = 0, 
        names.arg = barNaming(populations_nemo_rhine$V2), las = 2, cex.names = 1.5, 
        cex.axis = 1.5, cex.lab = 1.5, cex.main = 2)
    abline(v = b[vertLinePos(populations_nemo_rhine$V2)] - 0.5, lwd = 7, col = "black")
}
```

Overview of ADMIXTURE plots from K=2 to K=7 for A. nemorensis based on the de novo pipeline.

##### 4.6.3 A. sagittata reference-based

```
populations_all = read.table("/home/hditt/Arabis/samples_rad3/variants/populations_all.csv", 
    header = F, sep = ",", stringsAsFactors = F)
fam_sag_rhine = read.table("/home/hditt/Arabis/samples_rad3/variants/ArabisSagRhine_pseudo_0.95miss_reprem_loci_hetrem20_snps.recode.vcf.fam")
populations_sag_rhine = populations_all[which(populations_all$V1 %in% fam_sag_rhine$V1), 
    ]
populations_sag_rhine = populations_sag_rhine[order(populations_sag_rhine$V2), 
    ]
populations_sag_rhine$V2 = as.factor(populations_sag_rhine$V2)
levels(populations_sag_rhine$V2) = c("A-1", "B", "A-2", "D", "E", "F", "G", 
    "H", "I")
populations_sag_rhine$V2 = as.character(populations_sag_rhine$V2)

cvtable = read.table("/home/hditt/Arabis/samples_rad3/variants/admix_SagRhine_30052018/ArabisSagRhine_pseudo_0.95miss_reprem_loci_hetrem20_snps.recode.vcf_CV.log", 
    sep = ",", header = T)
cvtable = cvtable[order(cvtable$K), ]
plot(cvtable$K, cvtable$CV, type = "b", xlab = "K", ylab = "Cross-validation error")
```

```
COLORS_mx = c("#E6EE9C", "#81C784", "#26A69A", "#00BCD4", "#039BE5", "#1976D2", 
    "#4527A0")
for (i in 2:6) {
    admix_table = read.table(sub("LLLL", as.character(i), "/home/hditt/Arabis/samples_rad3/variants/admix_SagRhine_30052018/ArabisSagRhine_pseudo_0.95miss_reprem_loci_hetrem20_snps.recode.vcf.LLLL.Q.converted"))
    admix_table = admix_table[match(populations_sag_rhine$V1, fam_sag_rhine$V1), 
        ]
    # admix_table=admix_table[populations_sag_rhine$V2!='G_res',]
    b = barplot(t(as.matrix(admix_table)), col = COLORS_mx, main = paste("K=", 
        as.character(i), sep = ""), ylab = "Ancestry", border = "grey25", space = 0, 
        names.arg = barNaming(populations_sag_rhine$V2), las = 2, cex.names = 1.5, 
        cex.axis = 1.5, cex.lab = 1.5, cex.main = 2)
    abline(v = b[vertLinePos(populations_sag_rhine$V2)] - 0.5, lwd = 7, col = "black")
}
```

Overview of ADMIXTURE plots from K=2 to K=6 for A. sagittata based on the reference pipeline.

##### 4.6.4 A. sagittata de novo

```
fam_sag_rhine = read.table("/home/hditt/Arabis/samples_rad3/stacks/denovo_M6_n6_new/popstats_sag/populations.snps.vcf.fam")

cvtable = read.table("/home/hditt/Arabis/samples_rad3/stacks/denovo_M6_n6_new/popstats_sag/admix/populations.snps.vcf_CV.log", 
    sep = ",", header = T)
cvtable = cvtable[order(cvtable$K), ]

plot(cvtable$K, cvtable$CV, type = "b", xlab = "K", ylab = "Cross-validation error")
```

```
COLORS_mx = c("#F0A3FF", "#0075DC", "#FFCC99", "#94FFB5", "#9DCC00", "#C20088", 
    "#003380", "#FFA405", "#FFA8BB")
for (i in 2:6) {
    admix_table = read.table(sub("LLLL", as.character(i), "/home/hditt/Arabis/samples_rad3/stacks/denovo_M6_n6_new/popstats_sag/admix/populations.snps.vcf.LLLL.Q.converted"))
    admix_table = admix_table[match(populations_sag_rhine$V1, fam_sag_rhine$V1), 
        ]
    admix_table = admix_table[populations_sag_rhine$V2 != "G_res", ]
    b = barplot(t(as.matrix(admix_table)), col = COLORS_mx, main = paste("K=", 
        as.character(i), sep = ""), ylab = "Ancestry", border = "grey25", space = 0, 
        names.arg = barNaming(populations_sag_rhine$V2[populations_sag_rhine$V2 != 
            "G_res"]), las = 2, cex.names = 1.5, cex.axis = 1.5, cex.lab = 1.5, 
        cex.main = 2)
    abline(v = b[vertLinePos(populations_sag_rhine$V2[populations_sag_rhine$V2 != 
        "G_res"])] - 0.5, lwd = 7, col = "black")
}
```

Overview of ADMIXTURE plots from K=2 to K=6 for A. sagittata based on the de novo pipeline.

### 5 Map of populations

```
geodata = read.table("/home/hditt/Arabis/trip/Arabis_allSamples.csv", sep = ",", 
    header = T)
geodata_pop = aggregate(cbind(Latitude, Longitude) ~ Population + status, data = geodata, 
    FUN = mean)

myMap <- get_map(location = c(8.35, 49.82, 8.45, 49.92), maptype = "watercolor", 
    source = "stamen", crop = T, zoom = 14)
ggmap(myMap, darken = c(0.4, "white")) + geom_point(data = geodata_pop, aes(x = Longitude, 
    y = Latitude, col = status), pch = 20, size = 10, alpha = 0.7) + theme(legend.position = "bottom", 
    legend.title = element_blank(), legend.text = element_text(size = 22), text = element_text(size = 15), 
    axis.ticks.x = element_blank(), axis.text.x = element_blank(), axis.ticks.y = element_blank(), 
    axis.text.y = element_blank()) + xlab("") + ylab("") + scale_color_manual(values = COLORS[c(4, 
    8)]) + scalebar(dist = 2, dd2km = T, model = "WGS84", x.min = 8.35, x.max = 8.442, 
    y.min = 49.821, y.max = 49.92, st.bottom = F, st.size = 6, st.dist = 0.025)
```

Figure 1: Overview map of the populations. Populations are colored by type.

```
myMap_zoom <- get_map(location = c(8.391, 49.85855, 8.3935, 49.8605), maptype = "toner-background", 
    source = "stamen", crop = T, zoom = 17)
ggmap(myMap_zoom, darken = c(0.1, "#ff943d")) + geom_point(data = geodata_pop, 
    aes(x = Longitude, y = Latitude, col = status), pch = 20, size = 20, alpha = 0.7) + 
    theme(legend.position = "bottom", legend.title = element_blank(), legend.text = element_text(size = 22), 
        text = element_text(size = 15)) + xlab("") + ylab("") + scale_color_manual(values = COLORS[c(4, 
    8)]) + scalebar(dist = 0.05, dd2km = T, model = "WGS84", x.min = 8.391, 
    x.max = 8.3932, y.min = 49.8587, y.max = 49.8605, st.bottom = F, st.size = 7, 
    st.dist = 0.05)
```

Figure 1: Zoomed section of the map of the populations. Populations are colored by type.
