## Supplementary material for "Strengths and potential pitfalls of hay-transfer for ecological restoration revealed by RAD-seq analysis in floodplain *Arabis* species": Document S3

### ITS sequence alignment

Sequence alignment of reference ITS sequences of the *A. hirsuta* group and ITS sequences of nine samples.

Sample sequence names are colored by their assigned species based on the alignment (red - *A. nemorensis*, blue - *A. sagittata/A.hirsuta*).

Nucleotides are arranged in blocks of 10 bp and the total alignment length is 612 bp.

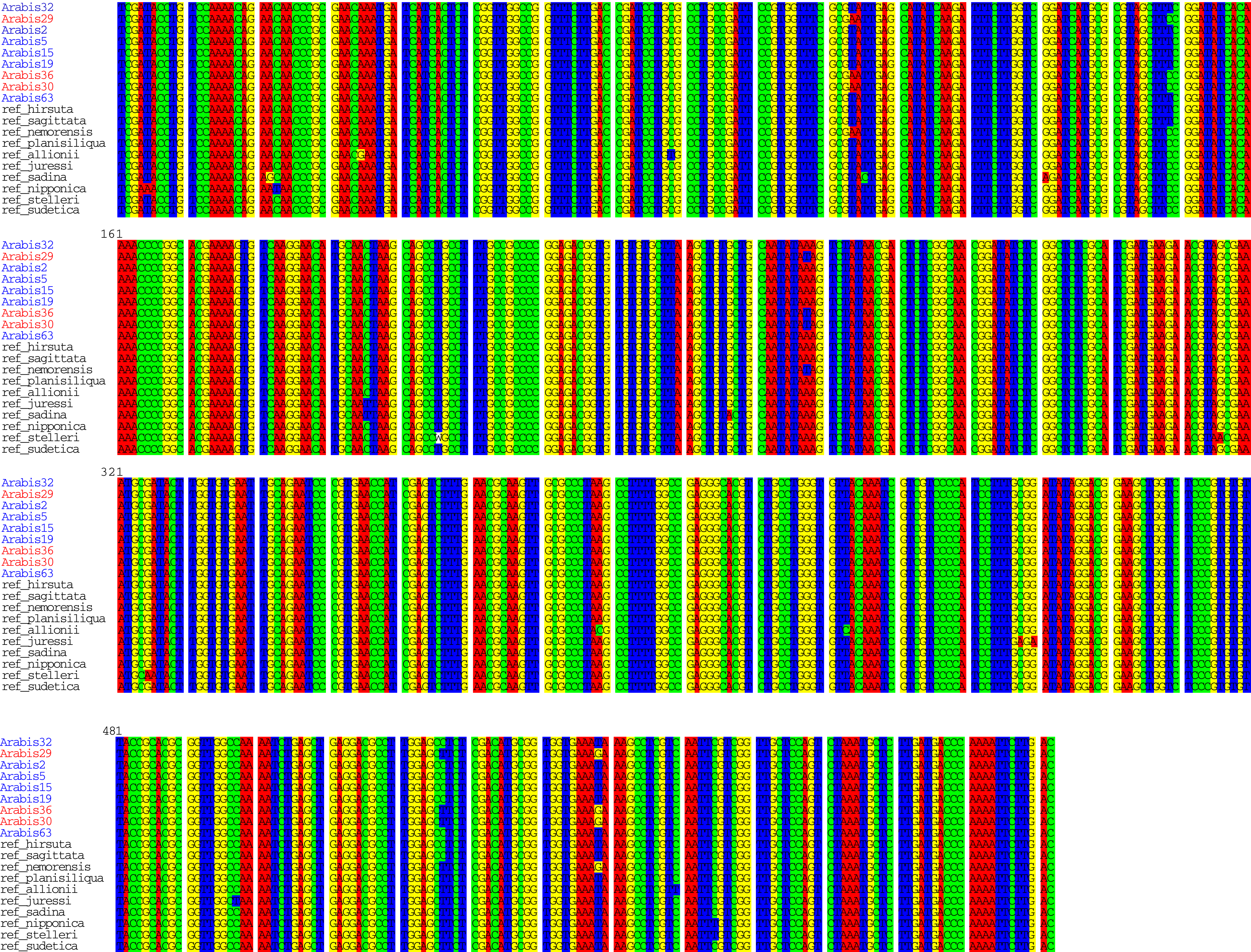
